# The BRD4S-LOXL2-MED1 interaction at the forefront of cell cycle transcriptional control in triple-negative breast cancer

**DOI:** 10.1101/2022.05.27.493725

**Authors:** Laura Pascual-Reguant, Tian V. Tian, Debayan Datta, Damiano Cianferoni, Savvas Kourtis, Antoni Gañez-Zapater, Chiara Cannatá, Queralt Serra-Camprubi, Lorena Espinar, Maria Guirola, Jessica Querol, Andrea Miró Canturri, Joaquin Arribas, Luis Serrano, Sandra Peiró, Sara Sdelci

## Abstract

Triple-negative breast cancer often develops resistance to single-agent treatments, which can be circumvented with targeted combinatorial approaches. Here, we demonstrate that the simultaneous inhibition of LOXL2 and BRD4 cooperate to reduce triple-negative breast cancer proliferation *in vitro* and *in vivo*. Mechanistically, we reveal that LOXL2 interacts in the nucleus with the short isoform of BRD4 and MED1 to control cell cycle progression at the gene expression level *via* sustaining the formation of BRD4-MED1 nuclear transcriptional foci. Indeed, the pharmacological or transcriptional repression of LOXL2 provokes downregulation of cell cycle gene expression, G1-S cell cycle arrest, and loss of BRD4-MED1 foci. Our results indicate that the BRD4S-LOXL2-MED1 interaction is fundamental for the proliferation of triple-negative breast cancer. Therefore, targeting such interaction holds potential for the development of novel triple-negative breast cancer therapies.

## Introduction

Bromodomain-containing protein 4 (BRD4) is an epigenetic reader well known in cancer for its role in the regulation of super-enhancers assembly (*1, 2*) and oncogenes transcriptional activation (*3–5*). Several BRD4 inhibitors, known as BET inhibitors (BETi), have been tested in multiple cancer models, and more than twenty clinical trials are currently ongoing to evaluate their anticancer efficacy (*6*). In breast cancer, the inhibition of BRD4 has shown promising preclinical results (*7*), sparking a particular enthusiasm for the treatment of the triple-negative breast cancer (TNBC) subtype, for which no targeted or efficient anticancer therapy has been developed so far (*8–10*). However, due to its heterogeneous and aggressive nature, TNBC commonly develops resistance to single-agent approaches (*11*), including BETi (*10*), which might be limited with the use of anticancer combinatorial approaches.

Lysyl Oxidase Like 2 (LOXL2) is a member of the lysyl oxidase family of copper-dependent amine oxidases (*12*) that catalyzes the oxidative deamination of peptidyl lysine residues. In the extracellular matrix, LOXL2 activity promotes collagen, elastin (*13*), and tropoelastin (*14*) crosslink, a phenomenon that is associated with the accumulation of extracellular matrix, fibrosis, and inflammation, which are all typical hallmarks of cancer (*15–17*). Intracellularly, LOXL2 can localize in the nucleus and promote the oxidation of nuclear proteins such as TAF10 (*18*) and Histone-3 (*19, 20*), leading to transcriptional repression and heterochromatinization, respectively.

Recently, it has been found that LOXL2 has a pivotal role in different solid cancers, including liver, pancreas, lung, and breast (*21–23*). LOXL2 inhibition efficiently reduces TNBC cell proliferation (*24*), sensitizes TNBC cells to common anticancer treatments such as DNA damaging agents (*25*), and inhibits the formation of TNBC distal metastasis (*21*). The mechanism by which LOXL2 can support cancer proliferation is not yet entirely understood and may vary among cancer types and subtypes.

In the present study, we addressed the question of whether the simultaneous inhibition of BRD4 and LOXL2 could be proposed as a novel strategy for the treatment of TNBC. Not only we discovered the two treatments synergize, but also that in the nucleus LOXL2 selectively interacts with the short isoform of BRD4 to promote the expression of cell cycle genes, thus controlling TNBC proliferation.

## Results

### The simultaneous inhibition of BRD4 and LOXL2 impairs TNBC proliferation

Given the fact that BRD4 and LOXL2 are both promising targets for the treatment of TNBC, we investigated whether the combination of their inhibition may cooperate in hindering TNBC proliferation. For this, we sought to perform a drug synergism analysis in three TNBC cell lines, MDA-MB-468, MDA-MB-231, and BT-549, which express different levels of BRD4 and LOXL2 (**Fig. 1A**). First, we verified that the LOXL2 inhibitor PXS5382 (*24*) efficiently reduced the nuclear catalytic activity of LOXL2 by checking the levels of oxidized Histone H3 (H3K4ox) in PXS5382-treated MDA-MB-231 cells, which were decreasing in a dose-dependent fashion (**fig. S1A**). MDA-MB-468, MDA-MB-231, and BT-549 cells were, therefore, treated either with PXS5382, or with the bromodomain inhibitor (*S*)-JQ1 (BETi) (*26*), or a combination of both small molecules for 96h. Both treatments given alone showed comparable results in the different cell lines when we assessed cell viability (**fig. S1B**). Interestingly, when the two treatments were given simultaneously we observed either an additive (in MDA-MB-468 and BT-549) or a synergistic (in MDA-MB-231) effect (**Fig. 1B and fig. S1C**).

**Fig. 1.**
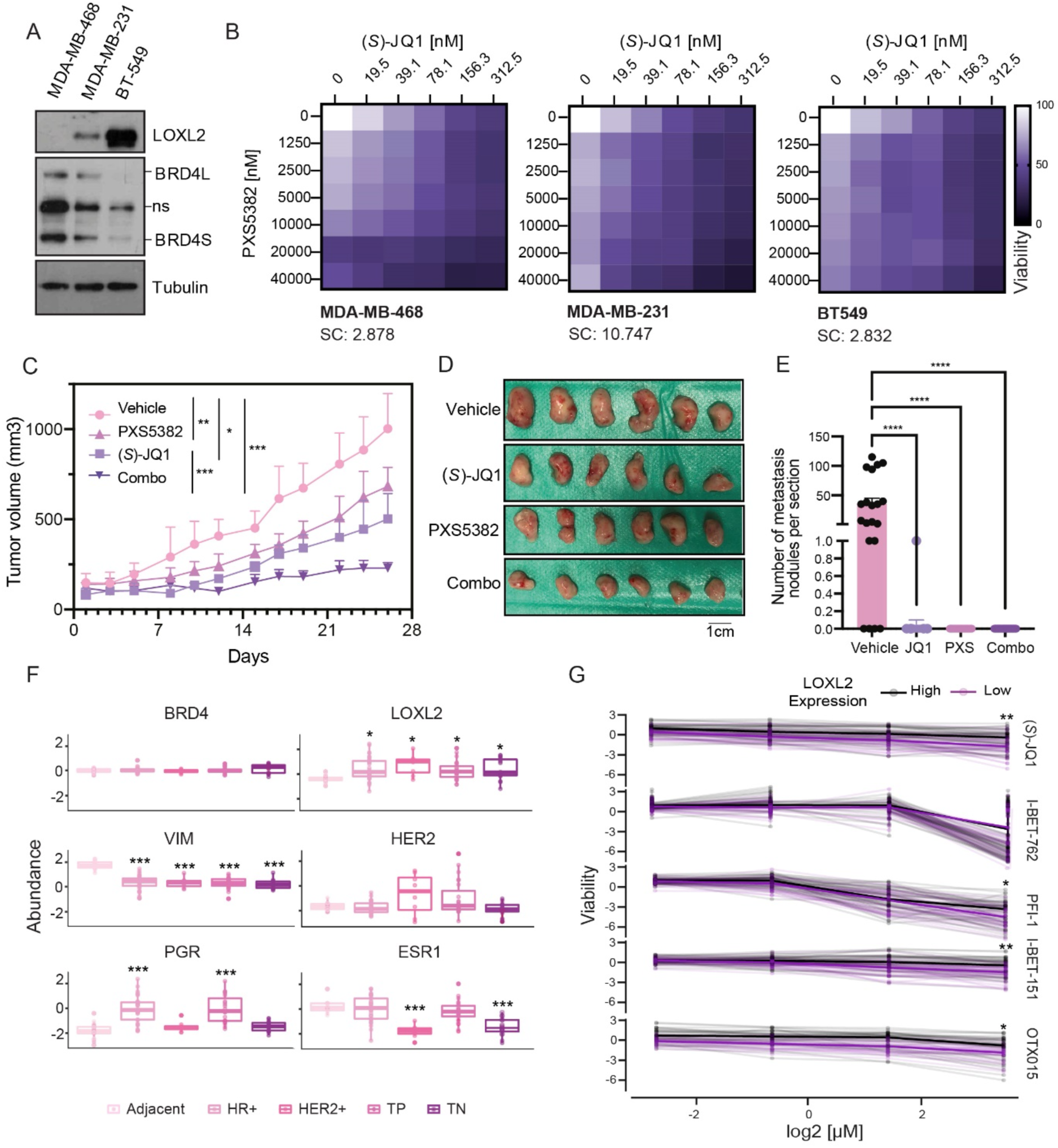
BRD4 and LOXL2 simultaneous inhibition impairs TNBC progression. (**A**) Representative Western blot analysis showing BRD4 and LOXL2 protein levels in the three different TNBC cell lines used. Tubulin is the loading control (ns: non-specific). Three biological replicates were performed. (**B**) Representative synergy matrixes showing cell viability measured with the MTT assay. The three TNBC cell lines used were treated with either PXS5382 or (S)-JQ1 alone or with their combination at the indicated concentration for 96h. SC indicates the synergy score for each cell line; a synergy score lower than 5 indicates the additive effect of the treatments, while a synergy score higher than 5 indicates synergism (*80*). Three biological replicates were analyzed for each cell line. (**C**) Tumor volumes from the MDA-MB-231 xenograft mice treated five times per week with 15mg kg^-1^ (*S*)-JQ1 and/or 2mg per pump PXS5382 during 26 days. 6 tumors per group (3 mice per group with one tumor on each side) are shown as average tumor volume, and standard deviations are shown as error bars. The asterisks indicate the significance at the endpoint (day 26) using a two-way ANOVA multiple comparisons with Tukey correction (\**P* < 0.05; ** *P* < 0.005; *** *P* < 0.001). (**D**) Images of the excised tumors at the end of the experiment (day 26). (**E**) Quantification of the number of metastasis nodules per mice lung section analyzed. The asterisks indicate the significance using a one-way ANOVA multiple comparisons with Dunnett correction (**** *P* < 0.0001). (**F**) Analysis of the CPTAC proteomics dataset showing the protein abundance of BRD4 and LOXL2 in different breast cancer subtypes from tumor samples classified by subtype using VIM, HER2, PGR, and ESR1 protein abundance. The asterisks indicate the significance of each cancer subtype against adjacent tissue using one-way ANOVA with Tukey’s posthoc (\**P* < 0.05; ** *P* < 0.005; *** *P* < 0.001; **** *P* < 0.0001). (**G**) Cell viability of high and low LOXL2-expressing CCLE cell lines (mRNA levels) treated with different BETi small molecules at the indicated concentrations. The asterisks indicate the significance at the highest concentration using an unpaired Student’s t-test with BH correction. (\**P* < 0.05; ** *P* < 0.005; *** *P* < 0.001; **** *P* < 0.0001).

As the combinatorial treatment in MDA-MB-231 cells showed a synergistic effect, we orthotopically implanted these cells into the mammary glands of immunodeficient mice (NOD SCID). When tumor volumes reached approximately 100mm3, MDA-MB-231 bearing mice were randomly divided into 4 groups, and treated with either vehicle, PXS5382, (*S*)-JQ1, or the combination of PXS5382 and (*S*)-JQ1 (combo). While PXS5382 or (*S*)-JQ1 alone were sufficient to delay tumor growth *in vivo*, the combo treatment showed clear superior effects (**Fig. 1, C and D**). Since MDA-MB-231 was derived from the pleural effusion of a TNBC patient, this cell line is known for its metastatic capacity when orthotopically implanted (*27*). Indeed, histopathologic analysis of lung sections from vehicle-treated mice showed tumor metastatic foci (**Fig. 1E and fig. S1D**). In contrast, such metastatic foci were observed neither in PXS5382-treated nor in combo-treated mice, confirming the potential anti-metastatic effect of LOXL2 inhibition (*21*). (*S*)-JQ1 treatment alone also showed great anti-metastatic potential, however, one metastatic nodule could still be found in the analyzed samples. Importantly, no significant toxicity was observed in combo-treated mice compared to vehicle-treated, PXS5382-treated, or (*S*)-JQ1-treated mice (**fig. S1E**).

We next tested whether the combination of PXS5382 and (*S*)-JQ1 was effective in a previously established TNBC PDX model (PDX-549) from EuroPDX (www.europdx.eu), in which we detected moderate protein levels of both BRD4 and LOXL2 (**fig. S1F**). Similar to what was observed in the MDA-MB-231 orthotopic xenograft model, also in the TNBC PDX model the combination of PXS5382 and (*S*)-JQ1 showed a superior anti-tumor effect to single-agent treatments (**fig. S1, G to I**). Again, no significant toxicity was observed in TNBC tumor-bearing mice treated with each inhibitor or their combination (**fig. S1J**). Taken together, these findings suggest that the simultaneous inhibition of LOXL2 and BRD4 can delay TNBC tumor growth *in vitro* and *in vivo*, and this novel strategy is promising for potential clinical application.

### LOXL2 does not control BRD4 expression

Since LOXL2 may induce chromatin compaction *via* H3K4 oxidation (*25*), while BRD4 works as a transcriptional activator, we presumed that the balance between the protein levels of BRD4 and LOXL2 might be the result of a transcriptional co-regulation. Cells expressing high levels of LOXL2 may downregulate BRD4 expression due to high chromatin compaction at the BRD4 promoter, resulting in low BRD4 protein levels. On the contrary, cells with low levels of LOXL2 would not repress BRD4 expression, showing high BRD4 levels. When overexpressing the Flag-tagged wild type form of LOXL2 (LOXL2wt) in the MDA-MB-468 cell line, where the endogenous LOXL2 is barely detectable, we could detect H3K4 oxidation at the BRD4 promoter followed by a mild reduction of BRD4 gene expression. The same effect was not achieved by overexpressing the catalytic dead form of LOXL2 (LOXL2m) (**fig. S2, A and B**). However, LOXL2wt overexpression was followed by minimal changes at the BRD4 protein levels (**fig. S2C**). Similarly, when downregulating LOXL2 in either MDA-MB-231 or BT-549 TNBC cell lines, which normally express medium to high levels of LOXL2, we could observe a decrease of H3K4 oxidation at the BRD4 promoter followed by a mild increase of BRD4 gene expression (**fig. S2, D to G**). Again, almost no changes were detected at the BRD4 protein levels (**fig. S2, H to I**). We wondered if by analyzing a broader cell cancer panel we could observe an inverse correlation between BRD4 and LOXL2 expression. The CCLE transcriptomics and proteomics datasets (*28*) did not show any correlation across lineages, between *LOXL2* and *BRD4* mRNA levels **(fig. S3A**) or protein abundances (**fig. S3B**). A similar result was observed when analyzing the TCGA proteomics data of human breast tumor samples (*29*) **(fig. S3C**). These data suggested that the synergism observed with the simultaneous inhibition of BRD4 and LOXL2 is not mediated by a transcriptional co-regulation mechanism driven by BRD4 and LOXL2 antithetic chromatin-associated roles.

We, therefore, asked whether breast cancer cells overall show increased protein levels of either LOXL2 or BRD4. By analyzing the CPTAC proteomics dataset (*30*), we stratified breast cancer samples into the different subtypes depending on the expression of common markers (VIM, HER2, PGR, and ESR1), and compared their LOXL2 and BRD4 protein levels with those of adjacent normal breast tissues. Interestingly, LOXL2 protein levels were significantly increased in every subtype of breast cancer, while BRD4 showed a mild increase only in the TNBC subtype (**Fig. 1F**). Additionally, when comparing the CCLE-associated BETi (*31*) sensitivity we could observe that cell lines expressing high levels of LOXL2 were less sensitive to BRD4 inhibition than LOXL2-low cell lines (**Fig. 1G**). A similar trend was shown when stratifying the cell lines based on LOXL2 protein levels (**fig. S3D**). These results suggested the possibility of a functional relationship between BRD4 and LOXL2, which we further investigated to unveil the molecular mechanism underlying the observed small molecules’ cooperation.

### LOXL2 interacts with the short isoform of BRD4 *via* its bromodomains but in an acetylation-independent manner

In the attempt of finding a possible functional connection between BRD4 and LOXL2, we wondered whether they may physically interact. To test this hypothesis, we extracted nuclei from MDA-MB-231 cells and performed a BRD4 pull-down (PD) experiment. Samples were then analyzed by western blot, and the results showed that LOXL2 was a BRD4 nuclear interactor (**Fig. 2A**). Given the fact that there are no efficient LOXL2 antibodies to perform endogenous LOXL2 PD, we carried out the complementary experiment by transiently overexpressing LOXL2wt in MDA-MB-231 and performing Flag PD instead. To our surprise, the results showed that LOXL2 selectively interacted with the short isoform of BRD4 (BRD4S) (**Fig. 2B**). The interaction was retained also when overexpressing LOXL2m, suggesting that the catalytic activity of LOXL2 is not required for its interaction with BRD4S (**Fig. 2B**). Comparable results were obtained when transfecting LOXL2wt or LOXL2m into HEK-293-T cells, in which LOXL2 expression is undetectable (**fig. S4A**).

**Fig. 2.**
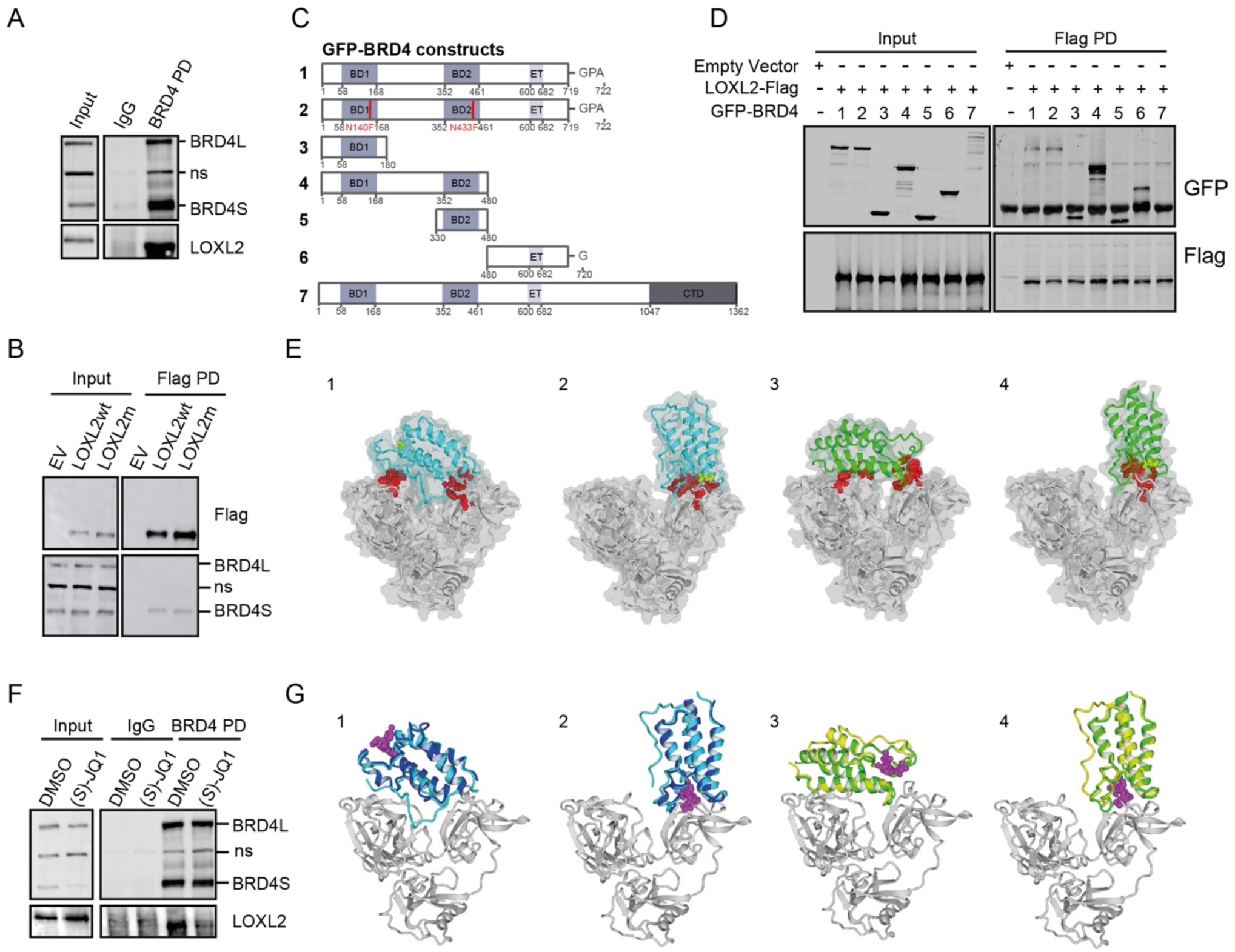
LOXL2 interacts with the short isoform of BRD4. (**A**) BRD4 PD in MDA-MB-231 cells using irrelevant IgG as a negative control. Precipitates were analyzed by Western blot with the indicated antibodies. Three biological replicates were performed (ns: non-specific). (**B**) Flag PD performed with MDA-MB-231 cells overexpressing either empty vector (EV), LOXL2-Flag wild type (LOXL2wt), or the catalytically dead form of LOXL2-Flag (LOXL2m). Precipitates were analyzed by Western blot with the indicated antibodies. Three biological replicates were performed (ns: non-specific). (**C**) Schematic representation of BRD4-GFP constructs used in Fig. 2c. (**D**) Flag PD of HEK293T cells overexpressing either empty vector (EV) or a combination of the indicated constructs. Precipitates were analyzed by Western blot with the indicated antibodies. Three biological replicates were performed. (**E**) Selected docking models, two for each of the BDs. 1 and 2 show BD1 structure (cyan) docked on LOXL2 (gray), corresponding to models 4uyd_zdock_10 and 4uyd_zdock_3, respectively (**table S3**). Figures 3 and 4 show models 2ouo_zdock_4 and 2ouo_zdock_5 (**table S3**) representing BD2 structures (green) docked into LOXL2 (gray). In red, the atomic representation of residues predicted to be fundamental for the modeled interaction (**table S3**). In yellow asparagines 140 (BD1) and 433 (BD2), respectively. The four models show their molecular volumes in light gray. (**F**) BRD4 PD in MDA-MB-231 cells treated either with DMSO or 5µM of (*S*)-JQ1 for 24h. IgGs were used as a negative control and the precipitates were analyzed by Western blot with the indicated antibodies (ns: non-specific). Three biological replicates were performed. (**G**) Superposition of selected docking models with BD1 (PDB: 3MXF) and BD2 (PDB: 3ONI) captured into crystallographic structures binding (*S*)-JQ1. (*S*)-JQ1 is shown in magenta and LOXL2 is shown in gray. Panel 1 shows BD1 (cyan) docking model 4uyd_zdock_10 superposed with 3MXF (blue), exhibiting compatibility of binding despite the presence of (*S*)-JQ1 into the AcK binding pocket. Panel 2 shows BD1 (cyan) docking model 4uyd_zdock_3 superposed with 3MXF (blue), exhibiting incompatibility of binding due to binding site competition of (*S*)-JQ1 and LOXL2. Panel 3 shows BD2 (green) docking model 2ouo_zdock_4 superposed with 3ONI (yellow), exhibiting compatibility of binding despite the presence of (*S*)-JQ1 into the AcK binding pocket. Panel 4 shows BD2 (green) docking model 2ouo_zdock_5 superposed with 3ONI (yellow), exhibiting incompatibility of binding due to binding site competition of (*S*)-JQ1 and LOXL2.

Since BRD4 binds to acetylated proteins (*9, 32–34*) *via* the Lysine acetylation (AcK) binding pocket of its bromodomains (BD1 and BD2), we wondered whether we could identify LOXL2 acetylated residues that would explain the interaction with BRD4S. The Phosphosite database (*35*) indicated that 4 different LOXL2 K-residues have been previously described as acetylated (K197, K209, K225 (*36*), and K248). Interestingly, one of them, K209, sits in a double-K H4-mimic similar motif (K209-K212) (**fig. S4B**), which is known to favor BRD4 binding to acetylated proteins like Histone-4 (*3, 37*) and the transcription factor Twist (*9*). However, we mutated the K209 residue to abrogate its acetylation (K→R), or the K209-K212 to mimic it (K→Q) and we did not observe any important change in LOXL2-BRD4S interaction (**fig. S4C**). These results indicated that likely the acetylation of these residues is not required for LOXL2-BRD4S interaction. Moreover, we overexpressed, in HEK293 cells, LOXL2wt together with the GFP tagged version of either BRD4S (1), BRD4S harboring inactivating mutations in BD1 and BD2 AcK binding pockets (N140F and N433F; 2), BRD4_BD1 (3), a fragment of BRD4 containing BD1 and BD2 (4), BRD4_BD2 (5), BRD4S-specific C-terminal domain (6), and BRD4 long isoform (BRD4L) (7) (**Fig. 2C)**. Flag-PD confirmed that LOXL2 interacted with BRD4S. BRD4S harboring mutations in the AcK binding pocket of both bromodomains still retained the ability to bind to LOXL2. In addition, both bromodomains either alone or together as well as the specific C-terminal domain of BRD4S interacted with LOXL2. Finally, although BRD4L was expressed at lower levels, we could not detect interaction with LOXL2, confirming results from Fig. 2b (**Fig. 2D**). Finally, we performed a docking analysis of BRD4_BD1/LOXL2 and BRD4_BD2/LOXL2, respectively. A collection of structures (**table S1**) from the Protein Data Bank (*38*) (PDB) and Interactome3d (*39*), and three independent software (ZDOCK (*40*), Autodock VINA (*41*), and ProteinFishing (*42*)) were used to propose binding models. Results were energetically minimized and ranked based on buried surface, FoldX (*43*) interaction energy, and FoldX (*43*) stability. A filtering step reduced the number of reliable docks to 7 (**table S2**). For those, we performed computational mutagenesis to exclude models incompatible with experimental data that showed that the mutations N140F of BD1 and N433F of BD2 still allowed LOXL2 binding (**Fig. 2D**). Such analysis reduced the candidate models to 4 (**Fig. 2E**). With these remaining models, we performed computational mutagenesis to highlight putative key-binding residues (**table S3**), which may be genetically perturbed to further dig down into the molecular basis of the interaction. In all the 4 proposed models, we observed that histidines 626 and 628 of LOXL2, which once mutated abrogate its catalytic activity (*19, 44*), did not participate in the interaction with BRD4_BDs (**fig. S4D**), corroborating the pull-down experiment presented in **Fig. 2B** showing that LOXL2m retained the interaction with BRD4S. Two of the 4 proposed binding models, were very similar between BD1 and BD2 and implicated the interaction of LOXL2 with the BD1/BD2 AcK binding pockets (models 2 and 4 in **Fig. 2E**). In these models, it is observable how the mutations N140F and N433F did not abrogate LOXL2 binding despite the strong involvement of the residues in the interaction (**fig. S4E**). For the remaining models (1 and 3) the interaction between BD1 or BD2 with LOXL2 did not involve the AcK binding pockets, making the N140F and N433F mutations irrelevant. We, therefore, reasoned that if the interaction between LOXL2 and BRD4_BDs would involve the AcK binding pocket most probably (*S*)-JQ1 treatment would abrogate it. Therefore, we treated MDA-MB-231 cells with (*S*)-JQ1 and performed a BRD4 PD experiment to investigate whether the treatment would impair BRD4-LOXL2 interaction. Results showed that the interaction is importantly reduced in presence of the treatment (**Fig. 2F**). In parallel, we observed that the presence of (*S*)-JQ1 invalidated the docking models 2 and 4 while did not perturb 1 and 3 (**Fig. 2G**), indicating that, most probably, the docking models 2 and 4 are the most accurate. Overall, these results indicated that the interaction between LOXL2 and BRD4S involves the AcK binding pocket of both BRD4S bromodomains, but that the residues N140 (BD1) and N433 (BD2), which are directly responsible for interacting with acetylated proteins, are dispensable. Additionally, the fact that the specific C-terminal domain of BRD4S mediates the interaction with LOXL2 may explain why BRD4L cannot engage in this interaction. Nonetheless, we cannot discard that the unstructured C-terminal domain of BRD4L may in addition function as a destabilizer.

### LOXL2 and BRD4S control the expression of DREAM target genes

Given the fact that LOXL2 and BRD4 were found to interact in the nucleus, we hypothesized that they may share transcriptional-associated functions. To dissect the possible transcriptional role executed by the BRD4S-LOXL2 interaction, we have performed and integrated ATAC-seq, RNA-seq, and BRD4-ChIP-seq comparing shControl (C) and shLOXL2 (KD) conditions in MDA-MB-231 cells (**fig. S5A**). The ATAC-seq experiment indicated that upon LOXL2 downregulation chromatin became much more relaxed, as expected due to its role in controlling chromatin compaction (**Fig. 3A**). Therefore, we initially expected that LOXL2 downregulation would induce gene expression changes towards upregulation. However, such a hypothesis was disproved since such changes were equally distributed between upregulated and downregulated genes (**Fig. 3B**). In addition, the increased accessibility observed upon LOXL2 downregulation was not followed by transcriptional activation (**Fig. 3C**). These data, therefore, showed that even though the loss of LOXL2 induced important chromatin changes, these were mostly not functional, or buffered against, as they did not correlate with gene expression changes. We then characterized the functional effect of LOXL2 downregulation on gene expression, independently of whether chromatin at that region was opening or not. When performing Gene Set Enrichment Analysis (GSEA), we observed that the downregulation of LOXL2 induced the upregulation of processes involved in cell morphology, secretion, membrane trafficking, and cell differentiation, with *cell-cell junction* being one of the most significant pathways (**fig. S5B**). These results are in agreement with the role of LOXL2 in reshaping the extracellular matrix (*45*) and regulating epithelial to mesenchymal transition (*46–48*), thus corroborating that the dataset we produced was of high quality. On the other hand, when we performed the same analysis on genes downregulated following LOXL2 KD we found that there was a significant enrichment of terms associated with cell cycle and, specifically, with DNA duplication (S-phase) and mitotic completion (M-phase) (**Fig. 3D**), suggesting a possible novel role of LOXL2 in controlling cell cycle progression. Of note, other downregulated GO terms, even if not directly, were still clearly associated with cell cycle regulation. Clear examples were *DNA conformational changes* that may relate to DNA duplication or chromatin condensation into chromosomes, and *organelle fission*, which refers to the process of mitochondrial fission that happens during the G2-M phase transition and is required to segregate mitochondria into the two daughter cells (**Fig. 3D**).

**Fig. 3.**
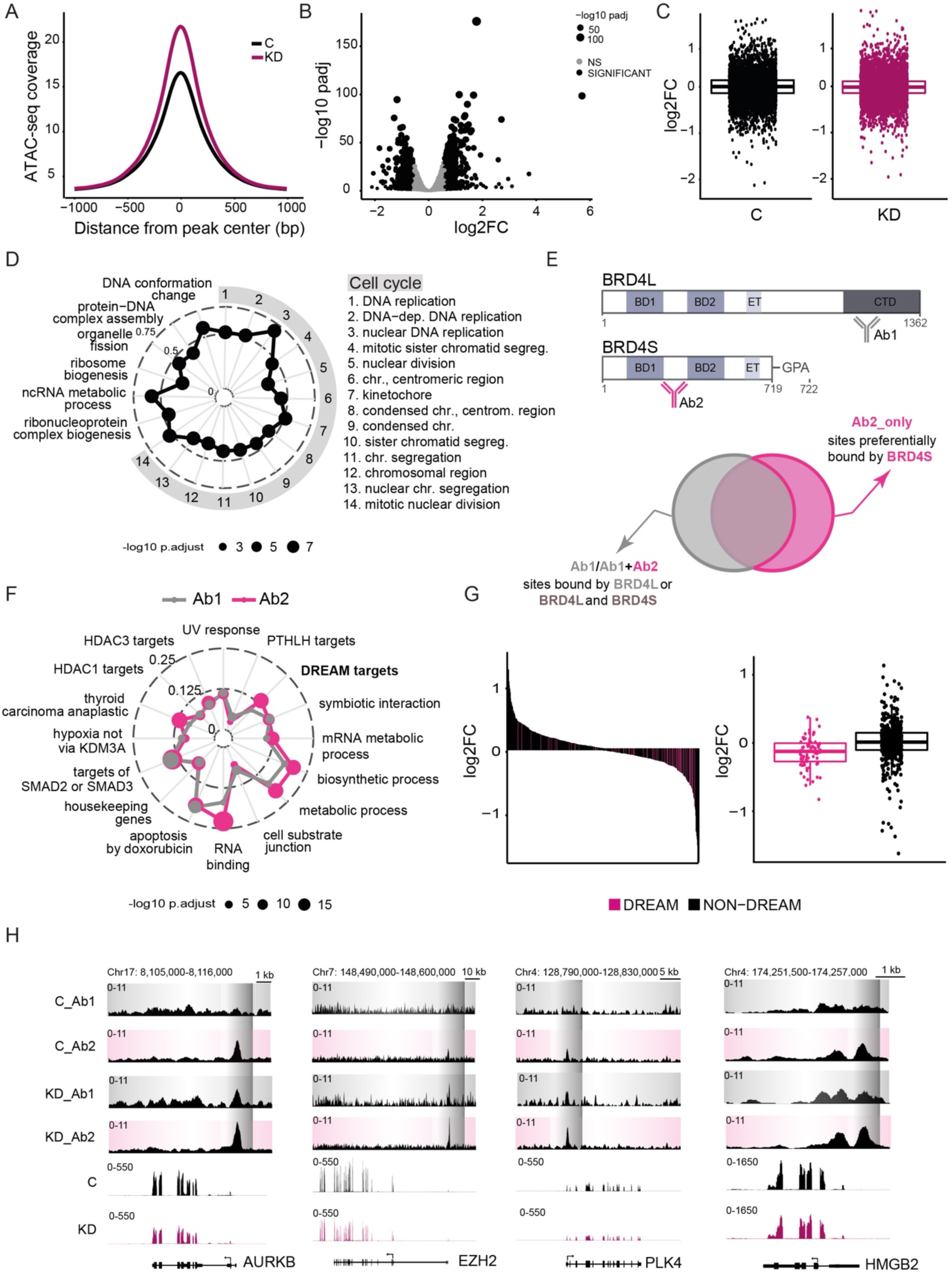
The interaction between LOXL2 and BRD4S regulates DREAM target gene expression. (**A**) ATAC-seq normalized coverage in MDA-MB-231 cells transduced with either shControl (C) or shLOXL2 (KD), represented as the distance from the center of all peaks (bp= base pair). (**B**) Volcano Plot representation of the differential expression of genes between C and KD conditions. Genes with adjusted p-values < 0.05 and abs(FC) > 1.5 are considered significant. (**C**) RNA-seq logFC for genes associated with ATAC-seq peaks in C and KD conditions which fall in promoter regions. (**D**) Gene Set Enrichment Analysis (GSEA) of the genes down-regulated upon KD. The top 20 categories are shown, with the size of the points proportional to the adjusted p-values and the distance from the center proportional to the gene ratio. (**E**) Schematic representation of BRD4L and BRD4S illustrating the Ab1 and Ab2 binding sites (top). Schematic representation of the ChIP-seq strategy used to identify BRD4S preferentially bound sites (bottom). (**F**) Overlap of promoter target GSs of Ab1 and Ab2 identified with the MSigDB collections. The size of the points is proportional to the adjusted p-values and the distance from the center is proportional to the gene ratio. The adjusted p-values are calculated independently for each overlap comparison (Ab1 and Ab2). (**G**) RNA-seq logFC for genes associated with the ChIP-seq peaks of Ab1 in C which fall in promoter regions. The logFC of the subset of these genes which are DREAM targets are plotted in pink. (**H**) Genome Browser tracks of four different DREAM target genes containing the following information (from top to bottom): Ab1 ChIP-seq profile, Ab2 ChIP-seq profile either in C or KD conditions, and RNA-seq signal in C and KD conditions.

We then asked how BRD4S and BRD4L bound across the genome in control conditions and whether their localization is affected by LOXL2 downregulation. We performed ChIP-sequencing (ChIP-seq) using two antibodies, one recognizing only BRD4L (Ab1) and one recognizing both isoforms (Ab2). We reasoned that the signal overlap between the two antibodies are chromatin regions bound either by BRD4L or BRD4L and S together, while the signal coming only from the Ab2 should correspond to BRD4S preferentially bound regions (**Fig. 3E**). We retrieved in total 2774 peaks for Ab1 and 3288 peaks for Ab2 and around 20% of the peaks were located at promoter regions (**fig. S5C**). With these identified promoters, we looked at overlaps with gene sets (GSs) in the Molecular Signatures Database (MSigDB) (*49*) to compare promoter regions differentially bound by the two antibodies. As expected, since both antibodies can bind to BRD4L, we observed an important functional overlap when comparing the top-10 identified GSs for each antibody (**fig. S5D**). When superimposing the results for Ab1 and Ab2, we observed that for some of the GSs Ab2 was showing a greater gene ratio and lower adjusted p-value (**Fig. 3F**), possibly suggesting that genes included in those pathways were stronger bound by BRD4S. Among them, the *Fisher_DREAM targets*, the *RNA binding*, and the *Rodrigues_thyroid carcinoma anaplastic up* GSs were the ones showing the greatest adjusted p-value and gene ratio differences. The *Fisher_DREAM targets* GS caught our attention because it comprises genes that regulate cell cycle progression (DREAM target genes) (*50*), which according to our RNA-seq, were deeply affected by LOXL2 downregulation (**Fig. 3D**). Interestingly, the majority of the DREAM target genes retrieved in our Ab2-ChIP-seq were downregulated in the LOXL2 KD condition (**Fig. 3G**), suggesting a functional interaction between BRD4S and LOXL2 transcriptionally controlling cell cycle progression.

We, therefore, tried to understand whether BRD4S and BRD4L relocalize differently on chromatin when LOXL2 is downregulated. For both antibodies, we observed a big increase in signal upon LOXL2 downregulation (**fig. S5E**), which agreed with the increased chromatin accessibility (**fig. S5F**). However, the increased BRD4 binding was not significantly followed by gene expression upregulation (**fig. S5G**), suggesting that it might just be a non-functional consequence of the increased chromatin accessibility. We then focused the analysis on DREAM target genes. We observed that in the absence of LOXL2 the promoters of the DREAM target genes were not anymore exclusively captured with Ab2, but with both antibodies, thereby indicating either a BRD4S-BRD4L co-binding or exclusive BRD4L binding (**Fig. 3H and fig. S5H**). We hypothesized that the increased signal observed with Ab1 and Ab2 in the KD condition may principally depend on BRD4L binding nonspecifically wherever chromatin becomes more accessible following the downregulation of LOXL2. Indeed, the overlap between the chromatin loci identified with the two antibodies considerably increased in the LOXL2 KD condition (**fig. S5I**). Altogether, these results suggested that LOXL2 and BRD4S may interact at the promoter of DREAM target genes to transcriptionally control cell cycle progression. Interestingly, the binding of BRD4L to DREAM target genes promoters in the absence of LOXL2 was followed by DREAM target genes’ transcriptional repression rather than upregulation.

### LOXL2 repression destabilizes the transcriptional control of early cell cycle genes

Given that 1) LOXL2 interacted with BRD4S, 2) that promoters of DREAM target genes were preferentially bound by BRD4S, and 3) that the downregulation of LOXL2 induced the transcriptional repression of DREAM target genes, we sought to investigate what was the underlying connection bringing BRD4S-LOXL2 interaction at the forefront of cell cycle transcriptional control. We first reasoned that if the downregulation of LOXL2 decreases the expression of DREAM target genes, it should also induce cell cycle arrest. We confirmed by qPCR that LOXL2 downregulation reduced the expression of a set of selected DREAM target genes in MDA-MB-231 cells (**fig. S6A**). When comparing the cell cycle profile of MDA-MB-231 cells treated with either C or KD, the latter was arrested in the G1 phase of the cell cycle (**Fig. 4A**). We then performed high-throughput (HT-) immunofluorescence (IF) using an antibody recognizing H3 serine 10 phosphorylation (H3S10p), a typical marker of mitotic entry, and quantified its signal in C and KD treated MDA-MB-231 cells. We observed that the percentage of positive H3S10p cells dramatically decreased when LOXL2 was downregulated, indicating that the absence of LOXL2 drastically hampered mitotic entry (**Fig. 4B and fig. S6B**). Thus, we wondered whether the role of LOXL2 in the regulation of cell cycle progression was dependent or independent of its catalytic activity. To answer this question, we treated MDA-MB-231 cells either with DMSO or PXS5382 and observed that LOXL2 inhibition considerably decreased the expression of DREAM target genes (**fig. S6C**), similarly to what was observed for the LOXL2 KD condition. This result indicated that the catalytic activity of LOXL2 is required for cell cycle transcriptional control. As a consequence, cells treated with PXS5382 were mostly arrested in the G1-S phase of the cell cycle, as observed by FACS (**Fig. 4C**). In addition, we performed a time-lapse experiment transducing MDA-MB-231 cells with vectors encoding for the protein SLBP, which accumulates in the nucleus during G1 and starts being degraded in G2 (*51*), tagged with the mTurquoise2 fluorophore, and the Histone H1, which allows nuclear identification, tagged with the Maroon1 fluorophore. When comparing DMSO *vs* PXS5382 treated cells, we observed that while DMSO-treated cells were able to progress into the cell cycle, first acquiring and then progressively losing the accumulation of nuclear SLBP-mTurquoise2, PXS5382-treated cells failed, retaining much longer the nuclear mTurquoise2 fluorescence, indicative of G1-S arrest (**Fig. 4D, fig. S6D and movie S1**).

**Fig. 4.**
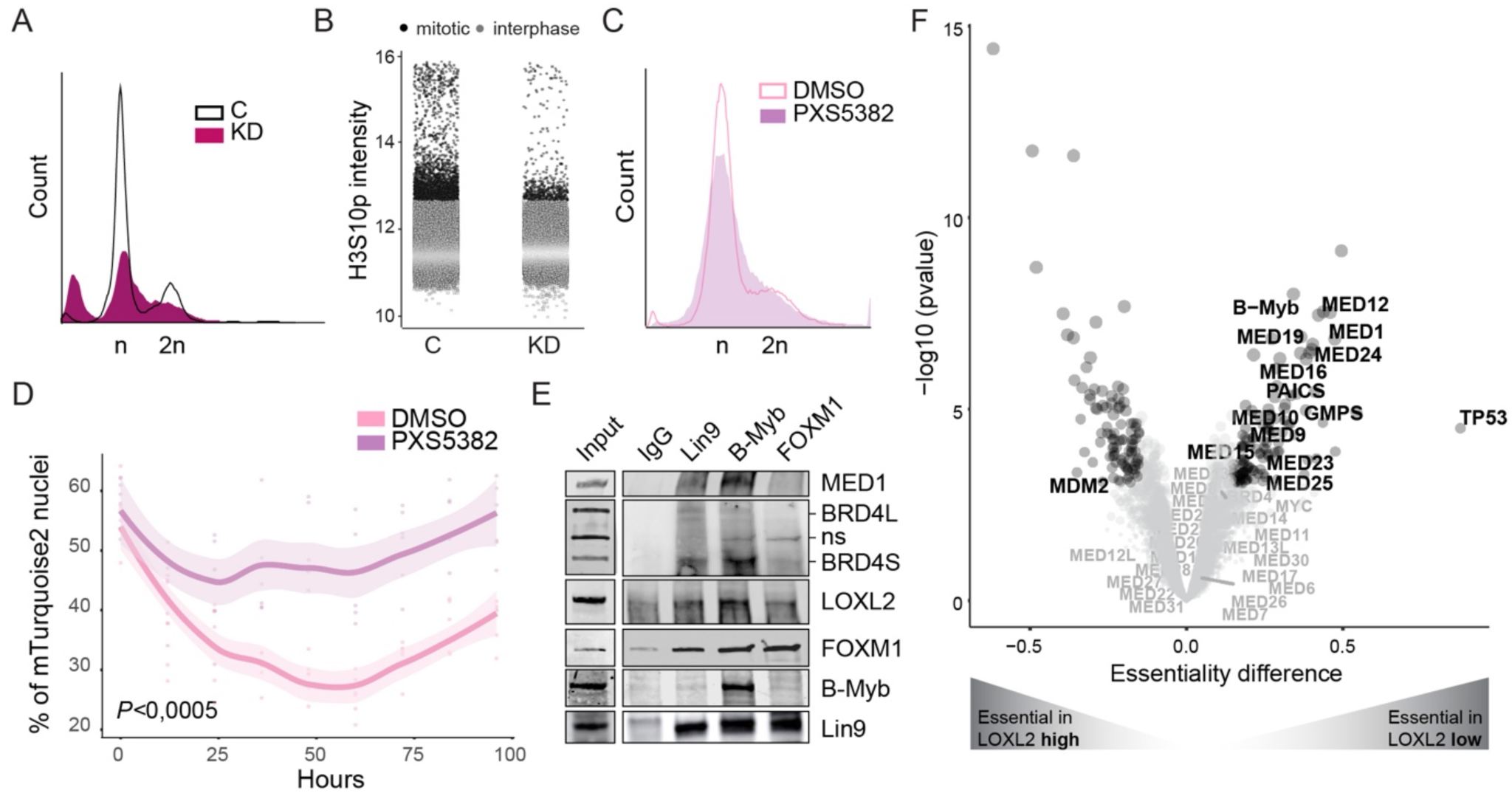
LOXL2 repression leads to G1-S arrest. (**A**) Representative cell cycle profile of MDA-MB-231 cells infected either with shControl (C) or shLOXL2 (KD). DNA content was analyzed by FACS following Hoechst staining. Three biological replicates were analyzed for each condition. (**B**) High-throughput immunofluorescence of H3S10p mitotic marker in C or KD transduced MDA-MB-231 cells. Mitotic cells (black dots) showed on average an H3S10p signal higher than the population median + 3S.D. Interphase cells are represented with gray dots. H3S10p intensity is represented as the normalized median. (**C**) Representative cell cycle profile of MDA-MB-231 cells treated either with DMSO or PXS5382 for 96h. DNA content was analyzed by FACS following Hoechst staining. Three biological replicates were analyzed for each condition. (**D**) MDA-MB-231 cells expressing SLBP-mTurquoise2 and H1-Maroon1 were treated with DMSO or PXS5382 for 96h. The percentage of mTurquoise2 nuclei in each well is shown, representing cells in G1-S. The difference between the PXS5382 and DMSO treated cells was significant (p-value <0.005) under a linear model comparing the percentage of mTurquoise2-positive cells, across time, in the two conditions. The estimated increase due to PXS5382 on the Area Under Curve (AUC), is 13.3 au. Quantification was performed every 12h. Six biological replicates were performed using at least 200 cells/replicate for the analysis. (**E**) PD of endogenous Lin9, B-Myb, and FOXM1 in MDA-MB-231 cells. Precipitates were analyzed by Western blot with the indicated antibodies. Irrelevant IgGs were used as a negative control (ns: non-specific). Three biological replicates were performed. (**F**) Differential gene essentiality between high and low LOXL2-expressing cell lines (CCLE) as calculated by analyzing the Achilles’ dataset. Cell lines with low LOXL2 expression are significantly more sensitive to depletion of genes represented in the right part of the X-axis as compared to cell lines with high LOXL2 expression, which are more sensitive to depletion of genes represented in the left part of the X-axis. Significance was determined using the Student’s t-test with BH multiple hypothesis correction. Significant threshold is based on adjusted p-value <0,05; black dots represent significant essentialities while gray dots are not significant. The dot size is proportional to the adjusted p-value of each gene.

The progression of the cell cycle relies on the fine-tuned transcriptional regulation of the DREAM target genes, which are divided into early and late cell cycle genes. The DREAM complex, which is composed of multiple subunits including the MuvB complex (LIN9, LIN37, LIN52, LIN54, and RBBP) and the Rb-like proteins p130, E2F4, and DP1, acts as a transcriptional repressor of cell cycle genes, which is the reason why they are called DREAM target genes. When cells start cycling (G1), the DREAM complex disassembles and only the MuvB complex remains associated with the promoter of DREAM target genes. Thus, the MuvB complex associates with E2F-class transcription factors that allow the transcriptional activation of G1-S transition-associated genes. On the other hand, the transcription of late cell cycle genes required for the S-G2 and G2-M transitions starts when the MuvB complex associates with the transcription factors B-MYB (S-G2) and later with FOXM1 (G2-M) (*52*). For this reason, early cell cycle genes often have E2F consensus sequences at their promoters, while late cell cycle gene promoters include B-MYB and FOXM1 consensus sequences (*50, 53*). Given the fact that when downregulating or inhibiting LOXL2 we observed a G1-S cell cycle arrest, we speculated that the BRD4S-LOXL2 interaction may be required for the transcription of early cell cycle genes. Supporting this hypothesis, when we conducted motif analysis on promoter regions recognized only with Ab2 we indeed observed enrichment in E2F consensus sequences (**table S4 and fig. S6E**).

We, therefore, wondered whether BRD4S and LOXL2 interact with the MuvB complex, B-Myb, and/or FOXM1. We performed Lin9 (MuvB subunit), b-MYB, and FOXM1 PD experiments using wild type MDA-MB-231 cells and showed that BRD4S and LOXL2 interacted with all the three factors, however, both proteins were stronger associated with Lin9 and B-MyB and milder with FOXM1 (**Fig. 4E**), again indicating that the BRD4S-LOXL2 interaction may be required for interphase cell cycle progression (G1-S-G2), rather than mitotic (G2-M). Additionally, we could barely detect BRD4L as either interacting with Lin9, b-MYB, or FOXM1, in agreement with the ChIP-seq results indicating that DREAM target gene promoters were preferentially bound by BRD4S (**Fig. 3F**). Interestingly, our PD experiments also showed that MED1 preferentially interacted with Lin9 and b-MyB, following the same pattern observed for BRD4S and LOXL2. Overall these data suggested that BRD4S, LOXL2, and MED1 all interacted with the MuvB complex and B-MyB to promote the transcription of early cell cycle genes. Interestingly, when performing an unbiased and orthogonal analysis exploring the Achilles dataset (*54*), we observed that b-MYB and several subunits of the Mediator complex (MED1, MED12, MED19, MED24, MED16, MED10, MED9, MED15, MED23, and MED25) scored as differentially top-essential in LOXL2 low-expressing cell lines compared to LOXL2 high-expressing ones. In addition, other BRD4 functionally related genes appeared as differentially essential in LOXL2 low-expressing cells, such as the de novo purine synthesis genes PAICS and GMPS (*55*). Finally, BRD4 itself, the BRD4 target oncogene MYC (*4, 5, 56, 57*), and the rest of the subunits of the Mediator complex showed a similar trend (**Fig. 4F and fig. S6F**), indicating that cells with reduced LOXL2 levels were more susceptible to the loss of BRD4 functional partners.

### LOXL2 catalytic activity is required for the stability of BRD4-MED1 transcriptional foci

It is very well known that the BRD4-Mediator interaction supports the formation of nuclear transcriptional foci (*58*) that decorate super-enhancer and boost the expression of target genes (*1, 59*). Given that BRD4 and MED1 colocalize at nuclear transcriptional foci (*58*), we investigated whether in our system we could detect such foci and if the downregulation of LOXL2 could impair their formation. We performed HT-IF in MDA-MB-231 cells either treated with C or KD, and immunostained for BRD4 or MED1. When quantifying the number of foci per area of the nucleus, we excitingly observed that in the KD condition there was a dramatic reduction of both BRD4 and MED1 foci (**fig. S7A**). Similarly, when comparing cells treated either with DMSO or PXS5382 we observed that the latter considerably reduced the number of BRD4 foci (**Fig. 5, A and B**) as well as the overlap between BRD4 and MED1 foci (**Fig. 5, C and B**). Consequently, the distance between a MED1 spot and the nearest BRD4 spot significantly increased (**Fig. 5D**). Given the fact that it has been recently discovered that BRD4S is crucial for the formation of BRD4-Mediator transcriptional foci (*60*), we wondered whether the downregulation of cell cycle genes observed with LOXL2 chemical or transcriptional inhibition could be the result of the destabilization of LOXL2-BRD4S-MED1 interaction. When performing a BRD4 PD we observed that the PXS5382 treatment did not impair the interaction between LOXL2 and BRD4, as expected from the docking results (**fig. S7B**). We, therefore, performed MED1 PD in DMSO and PXS5382 treated cells. Interestingly, we observed that in the DMSO condition MED1 was able to interact with LOXL2 but the PXS5382 treatment abolished this interaction (**Fig. 5E**). On the other hand, even though (*S*)-JQ1 treatment impaired the interaction between LOXL2 and BRD4 as previously observed (**Fig. 2F)**, it only mildly affected the interaction between BRD4 and MED1 (**Fig. 5F**). Therefore, the BRD4S-LOXL2-MED1 triangular interaction is only partially affected by solely inhibiting either BRD4 or LOXL2, thus explaining why the combinatorial treatment showed synergy in the tested TNBC models (**Fig. 5G**). Overall, these data revealed a novel mechanism by which the interaction between BRD4S and LOXL2 is required for the formation of BRD4-MED1 transcriptional foci and the gene expression regulation of early cell cycle genes (**Fig. 5H**).

**Fig. 5.**
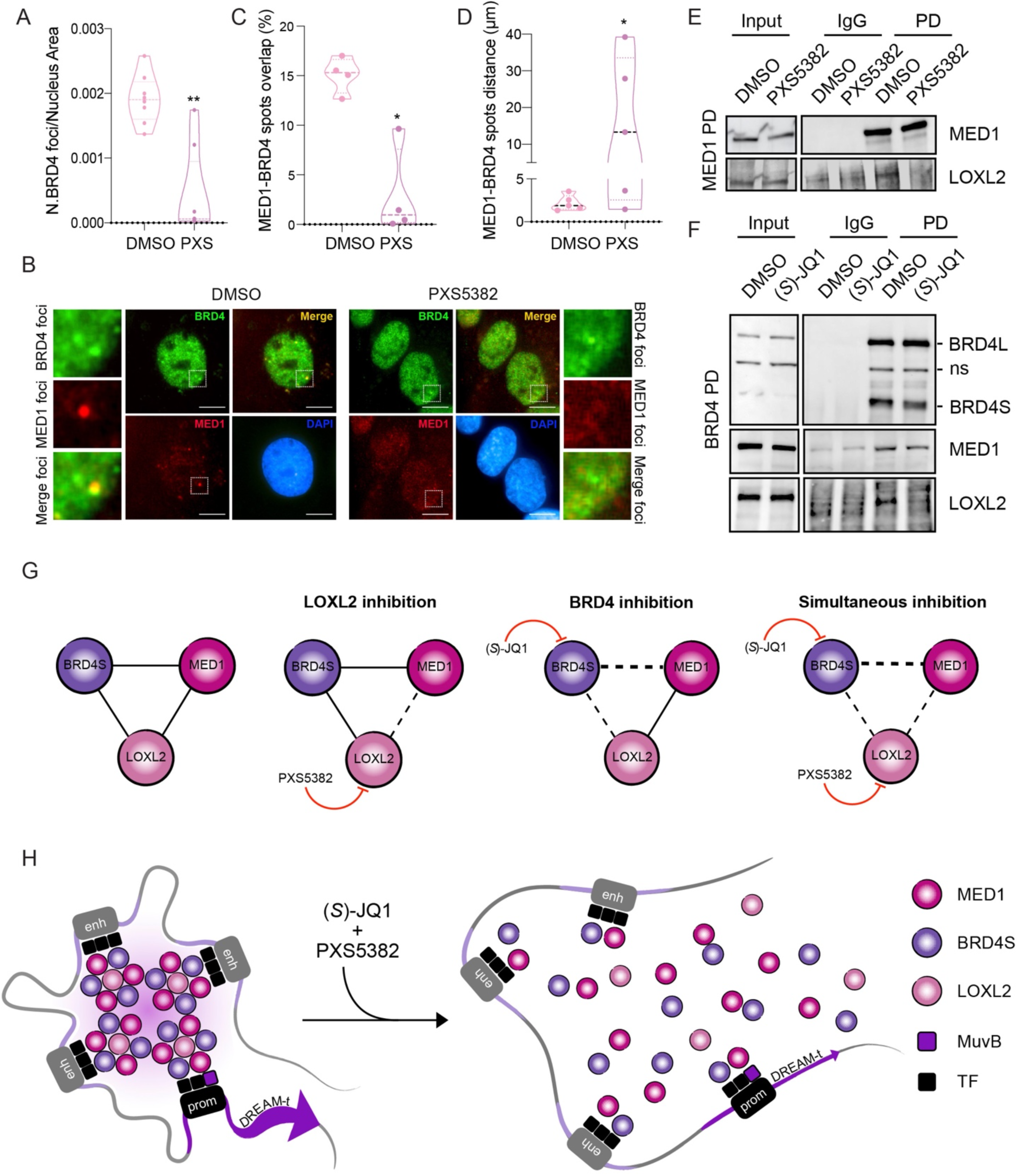
LOXL2 repression induces BRD4-MED1 transcriptional foci disintegration. (**A**) High-throughput immunofluorescence of BRD4 and MED1 in MDA-MB-231 cells treated with DMSO or PXS5382 for 96h. Quantification of BRD4 foci number corrected by the nucleus area is shown. Eight biological replicates were performed using at least 4000 nuclei/replicate for the analysis. The asterisks indicate the significance using a paired Student’s t-test (** *P* < 0.01). (**B**) Representative images of BRD4 (green) and MED1 (red) high-throughput immunofluorescence. DAPI (blue) was used as a nuclear marker. Scale bar; 5µm. Magnifications of representative foci are also depicted. (**C**) Quantification of the percentage of overlap between the area of each MED1 spot with that of BRD4 spots. Four biological replicates were performed using at least 4000 nuclei/replicate for the analysis. The asterisks indicate the significance using a paired Student’s t-test (* *P* < 0.05). (**D**) Quantification of the distance between each MED1 spot and its nearest BRD4 spot. Five biological replicates were performed using at least 4000 nuclei/replicate for the analysis. The asterisks indicate the significance using a paired Student’s t-test (* *P* < 0.05). (**E**) MED1 PD in MDA-MB-231 cells treated with DMSO or PXS5382 for 96h. Precipitates were analyzed by Western blot with the indicated antibodies. Irrelevant IgGs were used as a negative control (ns: non-specific). Three biological replicates were performed. (**F**) BRD4 PD in MDA-MB-231 cells treated with DMSO or (*S*)-JQ1 for 24h. Precipitates were analyzed by Western blot with the indicated antibodies. Irrelevant IgGs were used as a negative control (ns: non-specific). Three biological replicates were performed. (**G**) Schematic representation of the interaction between BRD4S-LOXL2-MED1 after inhibition of either BRD4, LOXL2, or the simultaneous inhibition of both proteins. (**H**) Schematic representation of the proposed molecular mechanism.

## Discussion

15% of breast cancers are classified as TNBC, they present a very aggressive phenotype and have an unfavorable prognosis. Given the fact that there is no targeted therapy to treat them, the standard regimen for TNBC treatment relies on the use of conventional chemotherapeutic agents. However, this strategy in most cases fails to arrest TNBC proliferation, whose molecular basis is largely unknown. In this study, we report an unprecedented mechanism controlling TNBC proliferation. While investigating whether the simultaneous inhibition of two promising TNBC targets -BRD4 and LOXL2-cooperates in arresting tumor proliferation, we discovered not only that such cooperation exists, but also that they are physical and functional partners. We showed that, in the nucleus, LOXL2 binds to the short isoform of BRD4 and that together they promote the expression of genes involved in the progression of the cell cycle interphase. For the very first time, we report LOXL2 as a transcriptional activator rather than a repressor, as previously described (*18*). We furthermore discovered that LOXL2 also interacts with MED1 and that LOXL2 catalytic activity favors the formation of BRD4-MED1 transcriptional foci. We hypothesize that the reduced formation of BRD4 and MED1 foci, which are known to decorate super-enhancer clusters, is the cause of the transcriptional repression of cell cycle genes observed either with LOXL2 downregulation or chemical inhibition. Interestingly, while LOXL2 has never been associated with cell cycle transcriptional control, BRD4 has been previously linked to the regulation of early cell cycle gene expression (*61, 62*), however, the molecular mechanism remained mostly unexplored. In addition, our work sheds further light on the divergent roles executed by different BRD4 isoforms, confirming the prooncogenic function of BRD4S (*63*). In this study, we provide in vivo data demonstrating that the simultaneous inhibition of LOXL2 and BRD4 can pave the way for the development of future TNBC anticancer approaches. Interestingly, the essentiality analysis that we conducted also revealed that LOXL2-low expressing cells are very sensitive to the loss of the oncosuppressor TP53, and conversely, they survive better if MDM2, which is required to induce TP53 proteasomal degradation, is absent (**Fig. 4F and fig. S6F**). This evidence is in line with previous results indicating that the loss of LOXL2 in TNBC enhances DNA damage and may indicate that cells with low levels of LOXL2 require a resilient mechanism to guarantee genome integrity and avoid apoptosis. Excitingly, also BRD4 has been multiple times associated with DNA damage response (*64, 65*), mechanistically being responsible either to insulate chromatin at the DNA damaged sites to allow repairing (*66*), or required to prevent the accumulation of R-loops and protecting against transcription–replication collision (*67*). The role of LOXL2 and BRD4 in DNA damage and their cooperation in controlling the transcriptional regulation of cell cycle progression described in this study may suggest that their simultaneous inhibition in tumor cells could act as a double-edged sword. In line with this, combining LOXL2 and BRD4 inhibition with DNA damaging agents could be expected to further improve the cancer treatment outcome. Although beyond the scope of this study, we believe that this approach may be suitable for the treatment of tumors where both BRD4 and LOXL2 are highly expressed.

## Materials and Methods

### Cell culture

HEK293T cells (ATCC No.: CRL-3216), MDA-MB-231 cells (ATCC No.: HTB-26), MDA-MB-468 (ATCC No.: HTB-132) and BT-549 (ATCC No.: HTB-122) were cultured in Dulbecco’s modified Eagle’s medium (Biowest; L0106-500) supplemented with 10% FBS (Gibco; 10270106) at 37ºC in 5% CO2. The lack of mycoplasma contamination was regularly tested every month.

### Plasmids

All the plasmids used in this study are listed below:

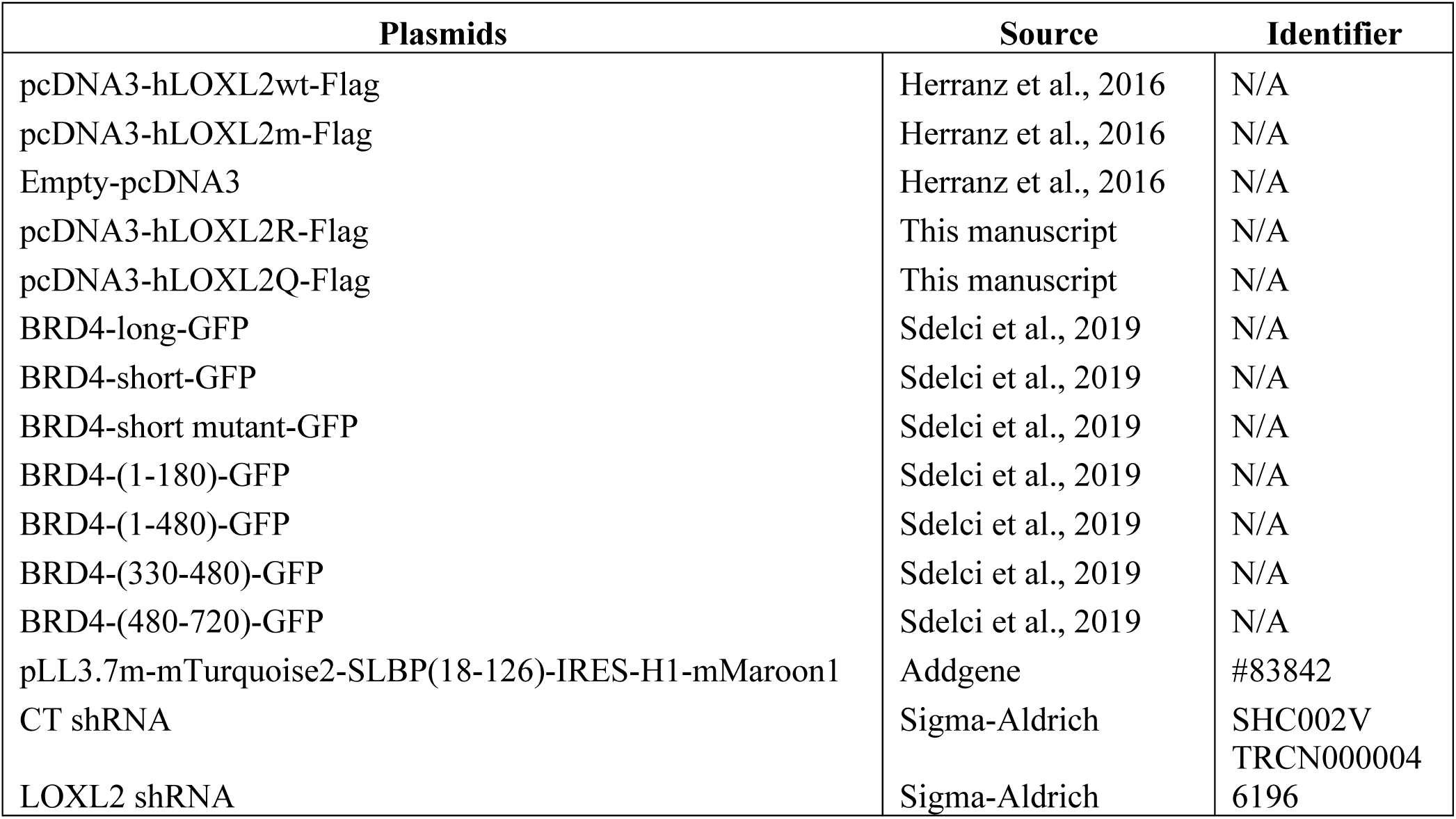

To obtain the human LOXL2 (hLOXL2) plasmid with point mutations in either the K209 or K209-K212, we designed and cloned a GeneBlock (IDT) containing the following mutations: K209R and K209Q-K212Q. 2µg of pcDNA3-hLOXL2 were cut with EcoRI and HindIII (New England Biolab) in cutsmart buffer for 2h and the cut plasmid was gel purified. We cloned each of the GeneBlocks containing the point mutations described above using Gibson reaction approach for 1 hour at 50ºC. Then, 3µl of Gibson reaction was transformed in DH5α *E*.*coli* and single colonies were grown in LB with ampicillin for further analysis. The primer 5’TACGACTCACTATAGGGAGACCCAAG3’ was used for Sanger sequencing selection of positive clones.

### Transfection

For overexpression assays, MDA-MB-231, MDA-MB-468, or HEK293T cells were seeded for 24h and transiently transfected with the indicated vectors using either polyethyleneimine polymer (Polysciences Inc; 23966-1) TransIT-X2 Dynamic Delivery System (Mirus Bio; MIR6004), following manufacturer’s instructions.

### Lentiviral infection

HEK293T cells were used for the production of lentiviral particles. Cells were grown to 70% confluence (day 0) and transfected by adding (dropwise) a mixture of 150 mM NaCl, DNA (50% of either Control shRNA [SHC002V] or LOXL2 shRNA [TRCN0000046196] vectors, 10% pCMV-VSVG, 30% pMDLg/pRRE, and 10% pRSV rev), and polyethyleneimine polymer (Polysciences Inc; 23966-1), which was pre-incubated for 15 min at room temperature. The transfection medium was replaced with fresh medium after 24h (day 1). On days 2 and 3, the cell-conditioned medium was filtered with a 0.45 μm filter unit (Merk Millipore; 051338) and stored at 4ºC. Viral particles were concentrated using Lenti-X Concentrator (Clonetech; 631232) following the manufacturer’s instructions, and virus aliquots were stored at –80ºC until use.

MDA-MB-231 and BT-549 cells were infected by adding the concentrated viral particles to their culture media. After 18h, the medium was replaced with fresh medium containing 2.5µg/ml of puromycin (Sigma-Aldrich; P8833). 48h after puromycin selection, the described experiments were performed.

### RNA extraction and qPCR

RNA was extracted using the PureLink RNA mini kit (Invitrogen) and converted into cDNA using the High Capacity RNA-to-DNA kit (Applied Biosystems) following the manufacturer’s instructions. qPCR was performed using Power SYBR Green PCR Master Mix (Applied Biosystems) in a 7900HT thermocycler (Applied Biosystems). For RNA-seq, samples were sequenced using the Illumina HiSeq 2500 system.

### Protein-protein docking analysis

Two structures covering BD1 and BD2 of BRD4 were retrieved from the Protein Data Bank (PDB) (*38*), and 45 models related to 6 observed protein-protein interactions involving BRD4 BDs were extracted from Interactome3d (*39*) (**table S1**). The first group of structures, together with the only available PDB model for LOXL2 (5ZE3), served to run docking on the ZDOCK server (*40*) and Autodock VINA (*41*). Models capturing BRD4 into interactions were used as input for ProteinFishing (*42*) together with the LOXL2 structure. Obtained models were later minimized through the YASARA structure minimization routine followed by FoldX *RepairPDB*. A reliability ranking by FoldX (*43*) interaction energy, stability, and the buried surface was generated (**table S1**) using FoldX *AnalyseComplex* and *Stability* commands, the buried surface was computed through the YASARA structure. The top 10 models for each BD were selected using the ranking mentioned above, models incompatible with LOXL2 AlphaFold (*68*) model AF-Q9Y4K0-F1 were excluded. Such a model predicts indeed the formation and packing of a domain implying residues from 1 to 318, unsolved in a LOXL2 PDB structure. Models with similar poses were grouped into clusters and further pruned by taking the top ranking one as representative of each cluster. For each of the six final models, residues involved in the interaction were determined using the FoldX *AnalyseComplex* command. FoldX *Pssm* command was used to predict the ΔΔG of mutating each of the interaction residues to Alanine. Mutations predicting a ΔΔG of interaction greater than 2.0 kcal/mol characterize the relative residue position as fundamental for the interaction. In the same way, ΔΔGs smaller than −2.0 kcal/mol should be considered invalidating for the relative model (**table S2**). Nevertheless, model *4uyd_zdock_3* was spared given the fact that unsatisfied residue D144 shows to possibly salt bridge with R447 of LOXL2. On the other hand, ProteinFishing model *O60885-O60885-EXP-5khm_5ze3_1* was excluded because of its lack of strong interactions to support the proposed binding pose (**table S2**). Asparagines 140 and 433 for BD1 and BD2 were mutated to phenylalanine and, in accordance with experimental data, models implying a loss of binding upon mutation were excluded (**table S3**).

### Cell cycle analysis

1 million MDA-MB-231 cells were trypsinized, resuspended in 1ml media, and stained with Hoechst 33342 (Invitrogen) 5 µg/ml final concentration for 30min at 37ºC in darkness. Stained cells were FACS analyzed using LSRFortessa and FlowJo V10 (BD Biosciences).

### Small molecule treatment and synergism analysis

Triple-negative breast cancer cell lines were seeded at day 1 in 96-well plates in triplicate: MDA-MB-468 (4000 cells/well), MDA-MB-231 (3000 cells/well), and BT-549 (3000 cells/well). The day after (day 2), cells were treated with the indicated concentrations of (S)-JQ1 or PXS5382, and their combinations. After 96h (day 6), MTT (3-(4,5-dimethylthiazol-2-yl)-2,5-diphenyltetrazolium bromide) assay was performed by adding 0.5 mg of MTT (Cat A2231, Panreac AppliChem) per mL of Dubelcco’s Modified Eagle Medium (DMEM) without Fetal Bovine Serum (FBS) for 3h at 37ºC to assess cell viability. The synergy score was calculated using the synergy finder 2.0 software (*69*).

### High-Throughput immunofluorescence

High-Throughput immunofluorescence analysis was performed with cells seeded on clear flat-bottom 96-well plates (Perkin Elmer), treated as described in the manuscript, and fixed with 4% paraformaldehyde for 10 minutes. Permeabilization and blocking were performed in PBS/3% bovine serum albumin (BSA)/0.1% Triton for 30 min. Thus, cells were incubated first with primary antibodies for 1h at room temperature (H3S10P 06-570 1:2000, BRD4 ab128874; 1:500, MED1 A300-793A; 1:500, Med1 LS-C290523; 1:200) and then with secondary antibodies (Alexa Fluor 488 Goat Anti-Rabbit and Alexa Fluor 555 Donkey Anti-Mouse, Thermo Fisher Scientific) for 30 min at room temperature and in the dark. Finally, cells were washed and incubated with DAPI (4,6-diamidino-2-phenylindole) for 5 min at room temperature and in the dark. Three PBS washing steps were done to remove the excess of antibodies and DAPI. Pictures were taken with the Operetta High Content Screening System (PerkinElmer), 63X or 40X objective, and non-confocal mode. Image quantification was done using the Harmony software, first identifying nuclei and then quantifying their properties (respectively, H3S10P intensity, BRD4 and MED1 foci, and BRD4/MED1 foci colocalization).

### Live cell imaging

MDA-MB-231 cells expressing mTurquoise2-SLBP(18-126) and H1-Maroon1 were seeded on clear flat-bottom 96-well plates (Perkin Elmer), treated with DMSO or PXS5382 40μM, and tracked for 96h with the Operetta High Content Screening System (PerkinElmer), 20X objective, and non-confocal mode. Image acquisition was done every 15 minutes and the quantification of mTurquoise2 and Maroon1 was done using the Harmony software.

### Western blot and pull-down experiments

Whole-cell extracts were obtained using SDS lysis buffer (2% SDS, 50 mM Tris-HCl, and 10% glycerol). Samples were mixed with 4x Laemmli sample buffer (Bio-Rad) and boiled at 95ºC for 5 min. Proteins were separated by SDS–polyacrylamide gel electrophoresis and the proteins were detected with the following antibodies: LOXL2 (Cell Signaling; 99680S), BRD4 (Abcam; ab128874), Tubulin (Sigma-Aldrich; T6557), and Flag (Sigma-Aldrich; F7425).

For the histone isolation experiment, cells were pelleted by centrifugation and washed with cold PBS. Pellets were resuspended by vortexing with lysis buffer (10 mM Tris, pH 6.5, 50 mM sodium bisulfite, 1% Triton X-100, 10 mM MgCl2, 8.6%sucrose, 10 mM sodium butyrate) and centrifuged at full speed for 15 seconds twice. The same procedure was repeated once with wash buffer (10 mM Tris [pH 7.4], 13 mM EDTA). Pellets were resuspended in 0.4 N H2S04 and left for 1h at 4ºC with occasional gentle mixing. After centrifuging at full speed for 5 min, supernatants were transferred to a new tube, and acetone was added (1:9). The mixture was left overnight at –20ºC and then centrifuged at full speed for 10 min. Pellets were air-dried for 5 min and resuspended in 30–100 μl of distilled water. Proteins were quantified using Bradford-Assay (Bio-rad; 5000006) and separated by SDS– polyacrylamide gel electrophoresis and the proteins were detected with the following antibodies; H3K4ox (previously generated 10.1038/s41388-019-0969-1) and H3 (Abcam; ab1791).

For the Flag pull-down experiment, MDA-MB-231 or HEK293T cells were transfected with pcDNA3-hLOXL2wt-Flag, pcDNA3-hLOXL2m-Flag, or an empty pcDNA3. After 48h, cells were washed twice with cold PBS and lysed in High Salt Lysis buffer (20 mM HEPES pH 7.4, 10% glycerol, 350 mM NaCl, 1 mM MgCl2, 0.5% Triton X-100) supplemented with protease inhibitors and incubated 30 min on ice the favor the lysis. Lysate samples were centrifuged at 13,000 rpm at 4ºC for 10 min. Balance buffer (20 mM HEPES pH 7.4, 1 mM MgCl2, and 10 mM KCl) was added to the resulting supernatant to reach a final NaCl concentration of 150 mM. Cell extracts were quantified using Pierce BCA Protein Assay Kit (Thermo Scientific; PIER23225) and 1mg was incubated with 40µl of Flag-M2 Affinity Agarose Gel (Sigma-Aldrich; A2220) for 4h at 4ºC and washed three times with wash buffer (20 mM HEPES pH 7.4, 1 mM MgCl2, 150 mM NaCl, 10% glycerol, 0.5% Triton X-100). Precipitated complexes were eluted with 2x Laemmli buffer. For endogenous pull-down experiments, cells were washed twice with cold PBS and lysed in Soft Salt Lysis buffer (50mM Tris-HCl pH 8.0, 10mM EDTA, 0.1% NP-40) supplemented with protease inhibitors for nuclear enrichment. After centrifugation at 3000rpm for 15min, nuclei were pelleted and lysed in High Salt Lysis buffer (20 mM HEPES pH 7.4, 10% glycerol, 350 mM NaCl, 1 mM MgCl2, 0.5% Triton X-100) supplemented with protease inhibitors and incubated 30 min on ice to favor the nuclei lysis. Lysate samples were centrifuged at 13,000 rpm at 4ºC for 10 min. Balance buffer (20 mM HEPES pH 7.4, 1 mM MgCl2, and 10 mM KCl) was added to the resulting supernatant to reach a final NaCl concentration of 150 mM. Nuclear extracts were quantified using Pierce BCA Protein Assay Kit (Thermo Scientific; PIER23225) and 1mg of nuclear extract was incubated with the indicated antibodies: 2.5ug of BRD4 (Abcam; ab128874), 5ug of MED1 (Bethyl Laboratories; A300-793A), 4ug of Lin9 (Proteintech; 17882-1-AP), 4ug of B-Myb (Proteintech; 18896-1-AP) 4ug of FOXM1 (Proteintech;13147-1-AP) O/N, followed by incubation with Protein A Dynabeads (Thermo Scientific; 10002D) for 1h at 4ºC. Complexes were washed three times with wash buffer and eluted with 2x Laemmli buffer. After pull-down experiments, proteins were separated by SDS–polyacrylamide gel electrophoresis, and the proteins were detected with the following antibodies: Flag (Sigma-Aldrich; F7425), GFP (Abcam; ab1218), BRD4 (Abcam; ab128874), Lin9 (Santa Cruz; sc-130571), FOXM1 (Santa Cruz; sc-376471), B-Myb (Santa Cruz; sc-81192), MED1 (Bethyl Laboratories; A300-793A), LOXL2 (Cell Signaling; 99680S)

### ATAC-seq sample preparation

Three biological replicates of ATAC-seq samples were prepared as previously described(*70*). Briefly, 50.000 MDA-MB-231 cells infected with shControl (C) or shLOXL2 (KD) were collected and treated with transposase Tn5 (Nextera Tn5 Transposase, Illumina Cat #FC-121-1030). DNA was purified using AMPure XP beads to remove big fragments (0.5x beads; >1kb) and small fragments (1.5x beads; <100bp). Samples were then amplified using NEBNexthigh-Fidelity 2x PCR Master Mix (New England Labs Cat #M0541) with primers containing a barcode to generate the libraries, as previously published (*71*). Each replicate was amplified with a combination of the forward primer and one of the reverse primers containing the adaptors listed in the primer’s table below. The number of cycles for library amplification was calculated as previously described (*70*). DNA was again purified using MinElute PCR Purification Kit (Qiagen) and samples were sequenced with Illumina HiSeq 2500.

### ATAC-seq analysis

Paired-end 50 bp reads were adaptor trimmed using TrimGalore (version 0.6.5). Trimmed reads were aligned to the hg19 genome (UCSC) using the BowTie 2 Aligner (version 2.4.2) (*72*) with parameters --very-sensitive -X 2000. Aligned reads were filtered using SAMtools (version 1.11) (*73*) to retain proper pairs with MapQ value >= 30. Read pairs aligning to ChrM were also discarded. Duplicate read pairs were removed with Picard (version 2.23.8). Read alignment was offset as previously described (*71*). Peaks were called using MACS2 (version 2.2.7.1) (*74*) with FDR < 0.01. Differentially accessible regions were determined using the DiffBind package (version 3.0.7) in R (version 4.0.2) with FDR < 0.005. The genomic annotation of the differential regions was performed with HOMER (version 4.11) (*75*). Normalized read coverage values were obtained using deepTools (version 3.5.0) (*76*). Coverage density heatmaps were generated using deepTools with the option reference-point and considering flanking regions of 1kb upstream and downstream from the center of the peaks.

### ChIP-qPCR and ChIP-seq sample preparation

10×10^6^ MDA-MB-231 cells infected with shControl (C) or shLOXL2 (KD) were crosslinked in suspension using 1% formaldehyde for 10 min at 37 °C. Crosslinking was stopped by adding glycine to a final concentration of 0.125 M for 5 min at room temperature. Cells were collected by centrifugation at 300g for 5min at 4ºC. Cell pellets were resuspended in SDS sonication lysis buffer (1M Tris-HCl pH 8, 10% SDS, 0.5M EDTA) supplemented with protease inhibitors. Extracts were sonicated to generate 200-600 bp DNA fragments, diluted 1:1,5 with equilibration buffer (10mM Tris, 233mM NaCl, 1.66% TritonX-100, 0.166% DOX, 1mM EDTA) supplemented with protease inhibitors and centrifuged at 14,000g for 10 min at 4ºC to pellet insoluble material. The primary antibody (5µg of H3K4ox (*25*), 5ug of Ab2; ab128874 or 15ug of Ab1; A301-985A100) or an irrelevant antibody (IgG) was added to the sample, and the mixture was incubated overnight with rotation at 4 °C (a 10% fraction was taken before adding the antibodies as an Input). Chromatin bound to the antibody was then immunoprecipitated using 25ul of protein A Dynabeads (Thermo Scientific; 10002D) for 2h with rotation at 4 °C. Precipitated samples were then washed two times with RIPA-LS (10mM Tris-HCl pH 8.0, 140mM NaCl, 1mM EDTA, 0.1% SDS, 0.1% DOX, 1% TritonX-100), two times with RIPA-HS (10mM Tris-HCl pH 8.0, 0.5M EDTA, 5M NaCl, 10% TritonX-100, 10% SDS, 10% DOX) and two times with RIPA-LiCl (10mM Tris-HCl pH 8.0, 1mM EDTA, 250mM LiCl, 0.5% NP-40, 0.5% DOX). Beads were resuspended in 48µl of ChIP elution buffer (10mM Tris-HCl pH 8.0, 5mM EDTA, 300mM NaCl, 0.4% SDS) and 2µl of Proteinase K and incubated at 55ºC for 1h and 65ºC O/N for de-crosslinking (inputs were also incubated for de-crosslinking).

DNA was purified with MinElute PCR purification kit (Qiagen) and eluted in nuclease-free water. Genomic regions of interest were detected by qPCR using the primers listed in the primers list. The NEBNext Ultra DNA Library Prep Kit for Illumina was used to prepare the libraries and samples were sequenced using Illumina HiSeq 2500 system.

### ChIP-seq Analysis

Single-end 50 bp reads were adaptor trimmed using TrimGalore (version 0.6.5).

Trimmed reads were aligned to the hg19 genome (UCSC) using the BowTie 2 Aligner (version 2.4.2) (*72*). Aligned reads were filtered using SAMtools (version 1.11) (*73*) to retain reads with MapQ value >= 30. Duplicate read pairs were removed with Picard (version 2.23.8). Peaks were called using MACS2 (version 2.2.7.1) (*74*) with FDR < 0.01. The genomic annotation of the differential peaks was performed with HOMER (version 4.11) (*75*). Normalized read coverage values were obtained using deepTools (version 3.5.0) (*76*). Motif Enrichment analysis was performed using HOMER (version 4.11) (*75*). Gene Ontology was performed using the clusterProfiler package (version 3.18.0) (*77*) in R (version 4.0.2). Gene overlaps with the MSigDB collections (version 7.4) were performed in http://www.gsea-msigdb.org/gsea/msigdb/annotate.jsp. Overlaps were calculated with C2 (curated gene sets), C5 (ontology gene sets), and C6 (oncogenic signature gene sets). Statistical significance of the distributions of ATAC-seq signals in ChIP-seq peaks was performed using a two-sample Kolmogorov-Smirnov test.

### RNA-seq sample preparation

Three biological replicates of RNA samples were prepared by using 1×10^6^ MDA-MB-231 cells infected with shControl RNA (C) or shLOXL2 RNA (KD). RNA was extracted using the PureLink RNA mini kit (Invitrogen) and sequenced in an Illumina HiSeq 2500.

### RNA-seq Analysis

50 bp single-end reads were aligned to the GRCh37.p13 Homo Sapiens reference genome using the STAR Aligner (version 2.7.6a) (*78*). Gene level counts were obtained by STAR using the --quantMode GeneCounts option, using gene annotations downloaded from Gencode (Release 19 GRCh37.p13). Differential expression analysis was performed in R (version 4.0.2) using the DESeq2 package (version 1.30.0) (*79*). Genes with adjusted p-value < 0.05 were considered differentially expressed. The lfcShrink function from DESeq2 was used for visualization purposes. Gene Set Enrichment Analysis (GSEA) was performed using the clusterProfiler package (version 3.18.0) (*77*) in R (version 4.0.2).

### Mouse xenograft studies

Mouse xenograft studies were performed using 6-week old NOD.CB17PrkdcSCID/J (NOD/SCID) female mice. For the generation of MDA-MB-231 orthotopic xenografts, one million low passage cells were diluted in Matrigel/PBS (v/v 1:1) and implanted into the number four fat pad of the mouse. In the case of patient-derived xenograft, low passage PDX549 (passage 5) was expanded and one 3mm diameter fragment was implanted into the mouse number four fat pad. After injection of cells and implantation of PDX549 into the fat pads, mice were sutured and kept in a clean cage with drinking water supplemented with Enrofloxacin (1,2mg/kg) for 2 weeks. Tumor xenografts were measured with calipers three times per week, and the tumor volumes were determined using the formula: (Length x width^2^)/2.

When the majority of tumor volumes reached 100mm3, mice were randomized into 4 groups: vehicle, (*S*)-JQ1, PXS5382, and Combo ((*S*)-JQ1 + PXS5382).

(*S*)-JQ1 was purchased from MedChemExpress (Cat HY-13030), and resuspended first in 10% DMSO, then added 40% PEG300, 5% Tween-80, and 45% sterilized saline buffer. 15mg/kg of the drug was administered by i.p. injection five times per week for a total of 4 weeks (cycles).

PXS5382 was provided by Dr. Wolfgang Jarlimek from Pharmaxis, Australia. The drug was resuspended in PBS buffer at 20mg/ml. 100 microliter of PXS5382-B6 solution was injected into an Alzet micro/osmotic pump (model 1004) according to manufacturer instructions (2mg/pump) and placed subcutaneously into the mice.

After 4 cycles of (*S*)-JQ1 treatment (Day 26), mice were euthanized and tumors were excised and measured. In addition, the lung of each mouse was dissected and fixed with 4% paraformaldehyde, then embedded in paraffin. Each lung sample was serially sectioned at 5µm of thickness, stained with H&E, and the 3rd, 5th, 15th, 25th, and 35th slides were used to count metastatic nodules.

### Primers

All primers used in this manuscript are listed below:

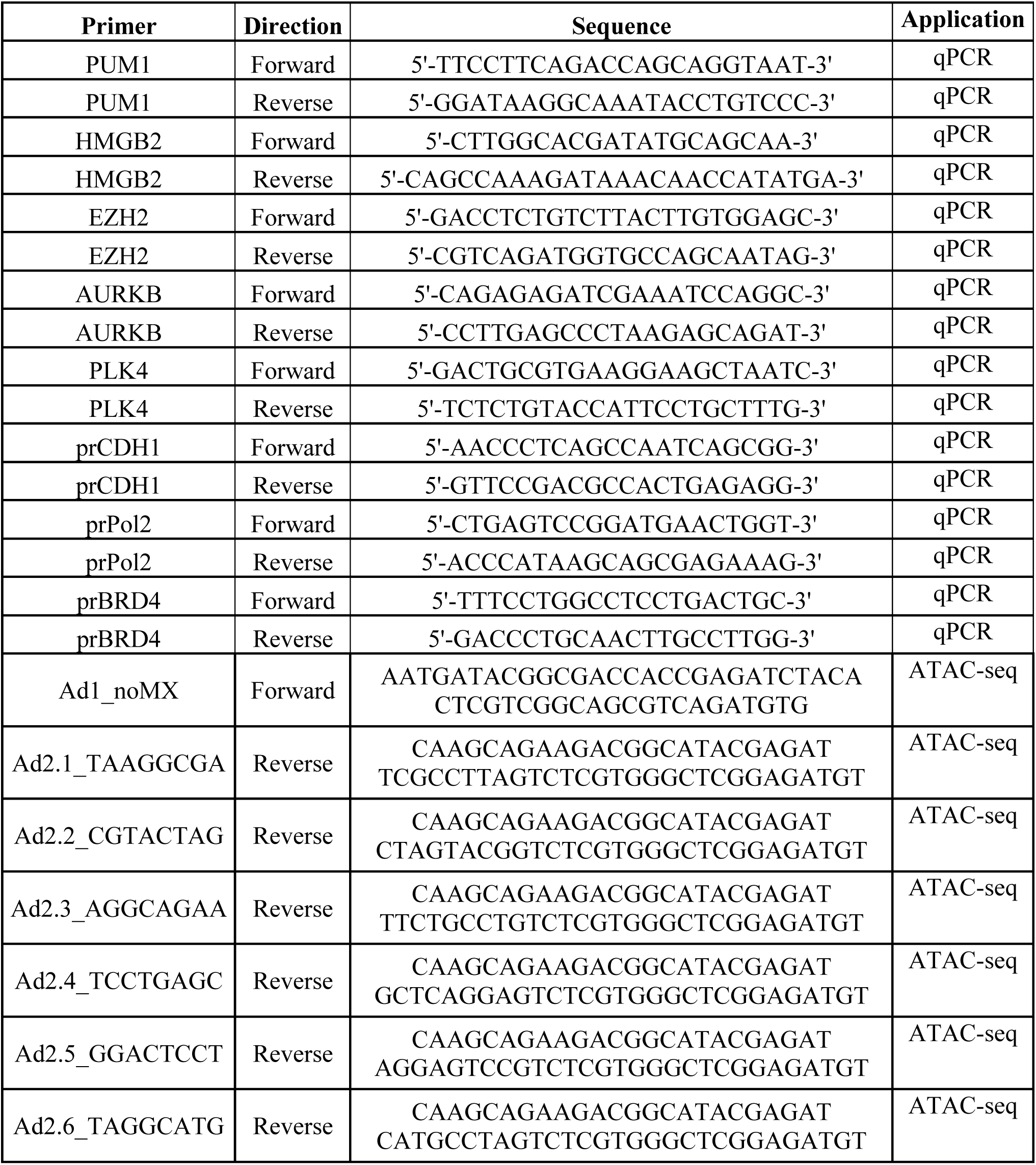

## Statistics

The statistical tests used are described within each figure legend.

### Data availability

Sequencing samples (raw data and processed files) are available at NCBI GEO under the accession number GSE198647

### Code availability

Differential essentiality analysis available on Github (https://github.com/Skourtis/LOXL2-BRD4).

### Contributions

S.S. S.P. and L.P.R. conceived the project and designed the study. L.P.R. performed all the cellular and molecular biology experiments shown in Fig. 2–4 and Supplementary Fig. 2-6. T.T. performed the in vitro and *in vivo* synergy analysis shown in Fig. 1 and Supplementary Fig. 1. D.D. performed the analysis of the ChIP-seq, ATAC-seq, and RNA-seq shown in Fig. 3 and Supplementary Fig. 4. D.C. performed the docking analysis shown in Fig. 2, Supplementary Fig. 4, and Supplementary Tables 2 and 3. S.K. performed the analysis of publicly available datasets shown in Fig. 1, Supplementary Fig. 1, Fig. 4, and Supplementary Fig. 6. A.G.Z. performed the time-lapse experiment shown in Fig. 4 and Supplementary Fig. 6. Q.S. contributed to the experiments shown in Supplementary Fig. 2. C.C. performed the PD experiment shown in Fig. 2c. L.E. performed the cloning of all the de novo vectors described in this manuscript. M.G. performed the PD experiment shown in Fig. 4f. J.Q. performed the Western blot shown in Fig.1e. L.S. contributed to the development of the docking analysis shown in Fig. 3 and Supplementary Tables 2 and 3. A.M.C and J.A. provided the PDX models. S.S. and L.P. wrote the manuscript with contributions from all the authors.

## Acknowledgment

We thank the CRG genomics unit, the CRG-UPF flow cytometry unit, and the VHIO mouse facility for their contribution. We thank Pharmaxis for the supply of PXS5382 LOXL2 inhibitors. S.S. is supported by the *Plan Estatal* de I+D+I (COMBAT PID2019-110598GA-I00), and the ERC Starting Grant (ERC-StG-852343-EPICAMENTE). L.P.R. is supported by the Juan de la Cierva-*Formación* fellowship (FJC2019-040598-I). T.T. is supported by *Plan Estatal* de I+D+I (PID2019-108008RJ-I00), AECC (INVES20036TIAN) and a Ramón y Cajal investigator contract (RYC2020-029098-I). D.C. is supported by the *la Caixa* Foundation PhD fellowship (ID 100010434; fellowship code LCF/BQ/DI19/11730061).

