## Supplementary Material for "The BRD4S-LOXL2-MED1 interaction at the forefront of cell cycle transcriptional control in triple-negative breast cancer"

**This PDF file includes:**

Figs. S1 to S7
Tables S1 to S4
Legends

**Other Supplementary Materials for this manuscript include the following:**

Movie S1

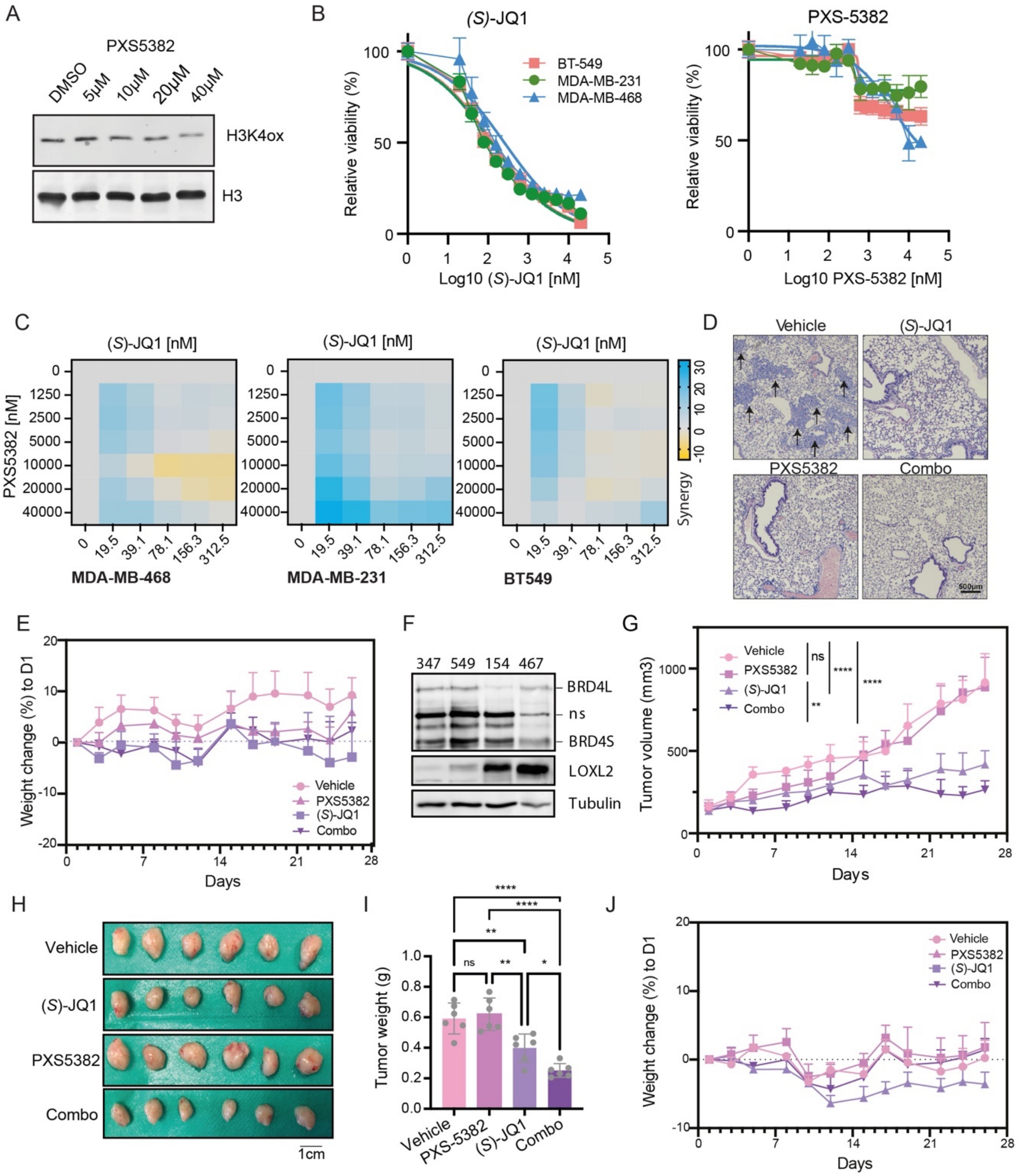
A
PXS5382
DMSO
5µM
10µM
20µM
40µM
H3K4ox
H3
B
(S)-JQ1
BT-549
MDA-MB-231
MDA-MB-468
Relative viability (%)
Log10 (S)-JQ1 [nM]
PXS-5382
Log10 PXS-5382 [nM]
C
(S)-JQ1 [nM]
PXS5382 [nM]
MDA-MB-468
MDA-MB-231
BT549
Synergy
D
Vehicle
(S)-JQ1
PXS5382
Combo
500µm
E
Weight change (%) to D1
Days
Vehicle
PXS5382
(S)-JQ1
Combo
F
347
549
154
467
BRD4L
ns
BRD4S
LOXL2
Tubulin
G
Tumor volume (mm3)
Days
Vehicle
PXS5382
(S)-JQ1
Combo
H
Vehicle
(S)-JQ1
PXS5382
Combo
1cm
I
Tumor weight (g)
Vehicle
PXS-5382
(S)-JQ1
Combo
J
Weight change (%) to D1
Days
Vehicle
PXS5382
(S)-JQ1
Combo

**Fig. S1**. **Effect of LOXL2 and BRD4 inhibition on TNBC proliferation and metastasis (A)** Western blot analysis of H3K4ox in MDA-MB-231 cells treated with DMSO or PXS5382 for 96h at the indicated concentrations; H3 was used as a loading control. Two biological replicates were performed. (**B)** Dose-response curves showing cell viability (MTT assay) of the TNBC cell lines tested treated either with (*S*)-JQ1 (left; 12 point dose-response, twofold dilutions starting from 20000nM) or PXS5382 (right; 12 point dose-response, twofold dilutions starting from 40000nM). (**C)** Representative matrixes showing the synergy score calculated with the cell viability data illustrated in Fig. 1b. (**D)** Representative H&E staining pictures of the mice lung sections from each group. Arrows indicate the presence of metastasis. **(E)** Mouse body weight changes from the MDA-MB-231 xenograft mice treated five times per week with 15mg $kg^{-1}$ (*S*)-JQ1 and/or 2mg per pump PXS5382 during 26 days. Weight changes are represented as the percentage to day 1 and standard deviations are shown as error bars. Bodyweight changes of less than 10% are considered tolerable. Statistical analysis was performed using two-way ANOVA multiple comparisons with Tukey correction for the whole experiment. **(F)** Representative Western blot showing LOXL2 and BRD4 protein levels of four different PDXs. Tubulin is the loading control (ns: non-specific). Three biological replicates were performed. **(G)** Tumor volumes from PDX549 mice treated five times per week with 15mg $kg^{-1}$ (*S*)-JQ1 and/or 2mg per pump PXS5382 during 26 days. 6 tumors per group (3 mice per group with one tumor on each side) are shown as average tumor volume, and standard deviations are shown as error bars. The asterisks indicate the significance at the endpoint (day 26) using a two-way ANOVA multiple comparisons with Tukey correction (*$P < 0.05$; ** $P < 0.005$; *** $P < 0.001$). (**H)** Images of the excised tumors at the end of the experiment (day 26). (**I)** Tumor weight from each group measured at the end of the experiment (day 26). Data are shown as average tumor volume weight and standard deviation are shown as error bars. The asterisks indicate the significance at the endpoint using a one-way ANOVA multiple comparisons with Tukey correction (*$P < 0.05$; ** $P < 0.005$; *** $P < 0.001$; **** $P < 0.0001$). (**J)** Mouse body weight changes from the PDX549 xenograft mice treated five times per week with 15mg $kg^{-1}$ (*S*)-JQ1 and/or 2mg per pump PXS5382 during 26 days. Weight changes are represented as the percentage to day 1 and standard deviations are shown as error bars. Body weight changes of less than 10% are considered tolerable. Statistical analysis was performed using two-way ANOVA multiple comparisons with Tukey correction for the whole experiment.

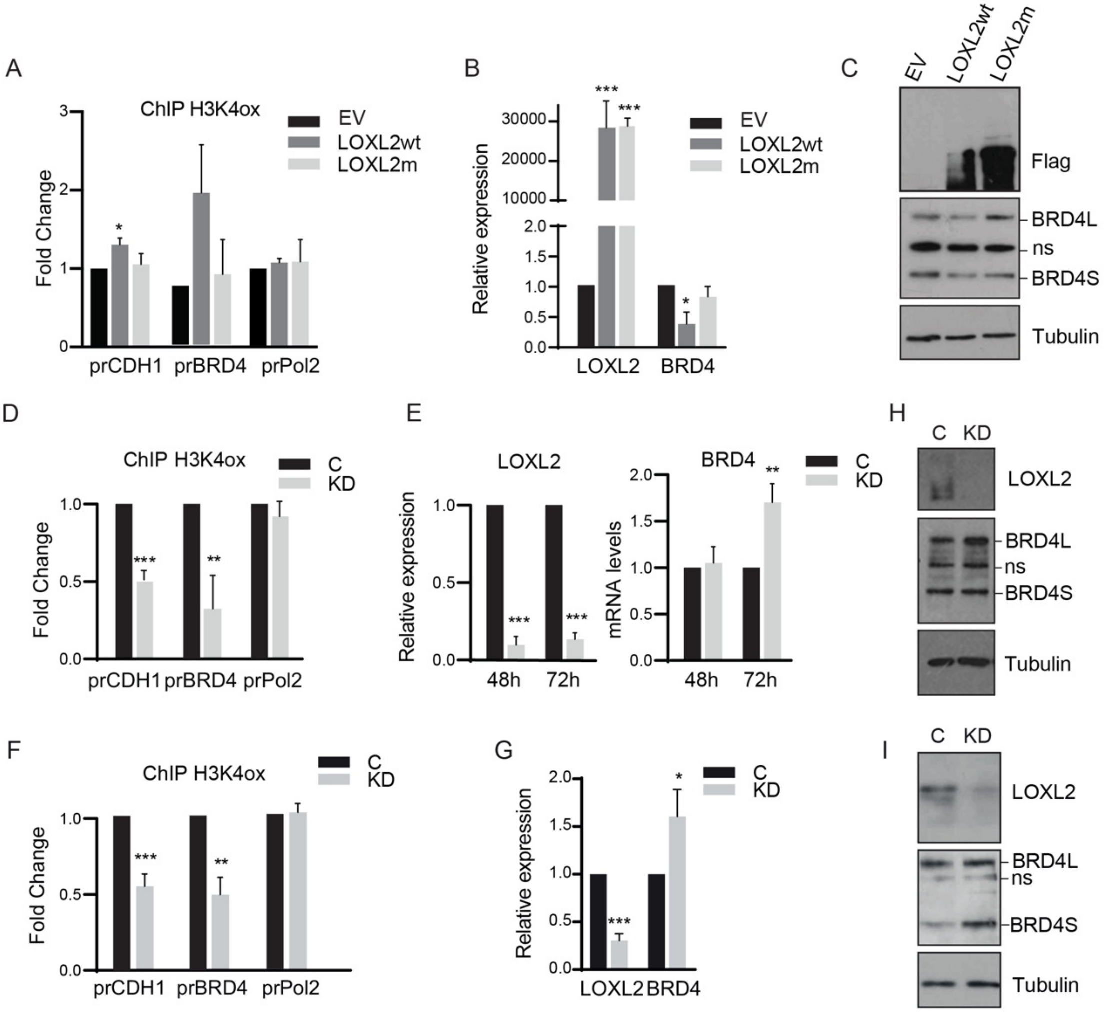
A
ChIP H3K4ox
EV
LOXL2wt
LOXL2m
Fold Change
prCDH1
prBRD4
prPol2
B
Relative expression
LOXL2
BRD4
C
EV
LOXL2wt
LOXL2m
Flag
BRD4L
ns
BRD4S
Tubulin
D
ChIP H3K4ox
C
KD
Fold Change
prCDH1
prBRD4
prPol2
E
LOXL2
BRD4
Relative expression
mRNA levels
48h
72h
H
C
KD
LOXL2
BRD4L
ns
BRD4S
Tubulin
F
ChIP H3K4ox
C
KD
Fold Change
prCDH1
prBRD4
prPol2
G
Relative expression
LOXL2
BRD4
I
C
KD
LOXL2
BRD4L
ns
BRD4S
Tubulin

**Fig. S2. Modulation of LOXL2 transcription does not induce changes on BRD4 protein levels**
(**A**) H3K4ox ChIP-qPCR of the BRD4 promoter (prBRD4) in MDA-MB-468 cells overexpressing either an empty vector (EV), LOXL2-Flag wild type (LOXL2wt), or the catalytically dead form of LOXL2-Flag (LOXL2m). E-cadherin promoter (prCDH1) and RNA Pol 2 promoter (prPol2) were used as positive and negative controls, respectively. Data from qPCR were normalized to the input and represented as the fold-change relative to EV condition, which was set as 1. Data are shown as the mean of three independent biological replicates and standard deviation as error bars. The asterisks indicate the significance using a one-way ANOVA multiple comparisons with Tukey correction (* $P < 0.05$). (**B**) Real-time RT-PCR (q-PCR) showing the changes in mRNA expression of LOXL2 and BRD4 in MDA-MB-468 cells overexpressing EV, LOXL2wt, or LOXL2m. Gene expression was normalized against an endogenous control (Pumilio homolog 1) and represented as the expression relative to the EV condition, which was set as 1. The asterisks indicate the significance using a one-way ANOVA multiple comparisons with Tukey correction (* $P < 0.05$). (**C**) Western blot of MDA-MB-468 cells overexpressing EV, LOXL2wt, or LOXL2m showing BRD4 and LOXL2 protein levels; Tubulin was used as a loading control and Flag as a transfection control (ns: non-specific). (**D**) H3K4ox ChIP-qPCR of the BRD4 promoter (prBRD4) in MDA-MB-231 cells treated with a shControl (C) or shLOXL2 (KD). E-cadherin promoter (prCDH1) and RNA Pol 2 promoter (prPol2) were used as positive and negative controls, respectively. Data from qPCR were normalized to the input and represented as the fold-change relative to the C condition, which was set as 1. The asterisks indicate the significance using a paired Student's t-test (*$P <$ 0.05; ** $P < 0.005$; *** $P < 0.001$). (**E**) Real-time quantitative RT-PCR (qRT-PCR) showing the changes in mRNA expression of LOXL2 and BRD4 in C and KD MDA-MB-231 cells. Gene expression was normalized against an endogenous control (Pumilio homolog 1) and represented as the expression relative to the EV condition, which was set as 1. The asterisks indicate the significance using a paired Student's t-test (*$P < 0.05$; ** $P < 0.005$; *** $P < 0.001$). (**F**) H3K4ox ChIP-qPCR of the BRD4 promoter (prBRD4) in BT-549 cells infected with a shControl RNA (C) or shLOXL2 RNA (KD). E-cadherin promoter (prCDH1) and RNA Pol 2 promoter (prPol2) were used as positive and negative controls, respectively. Data from qPCR were normalized to the input and represented as the fold-change relative to the C condition, which was set as 1. The asterisks indicate the significance using a paired Student's t-test (*$P < 0.05$; ** $P < 0.005$; *** $P < 0.001$). (**G**) Real-time quantitative RT-PCR (qRT-PCR) showing the changes in mRNA expression of

LOXL2 and BRD4 in C and KD BT-549 cells. Gene expression was normalized against an endogenous control (Pumilio homolog 1) and represented as the expression relative to EV condition, which was set as 1. The asterisks indicate the significance using a paired Student's t-test (*$P < 0.05$; ** $P < 0.005$; *** $P < 0.001$). (**H**) Western blot of shControl and shLOXL2 treated MDA-MB-231 cells showing BRD4 and LOXL2 protein levels; Tubulin was used as a loading control (ns: non-specific). (**I**) Western blot of C and KD MDA-MB-231 cells showing the differences in BRD4 and LOXL2 protein levels; tubulin was used as a loading control (ns: non-specific).

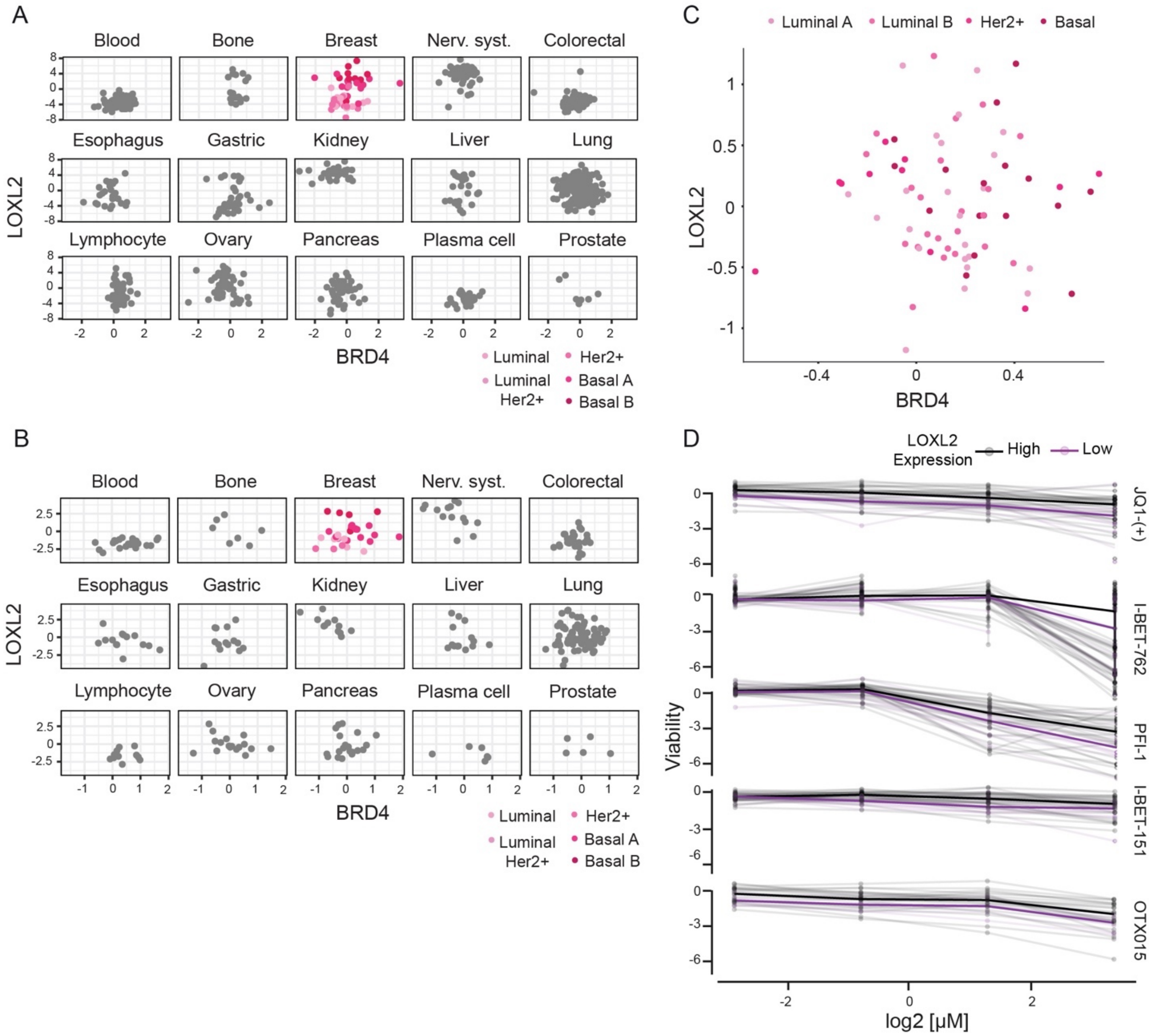
A
Blood
Bone
Breast
Nerv. syst.
Colorectal
Esophagus
Gastric
Kidney
Liver
Lung
Lymphocyte
Ovary
Pancreas
Plasma cell
Prostate
LOXL2
BRD4
Luminal
Luminal Her2+
Her2+
Basal A
Basal B
B
C
Luminal A
Luminal B
Her2+
Basal
D
LOXL2 Expression
High
Low
Viability
JQ1-(+)
I-BET-762
PFI-1
I-BET-151
OTX015
log2 [μM]

**Fig. S3. LOXL2 and BRD4 levels are not inversely correlated across tumors** (**A and B**) Correlation between LOXL2 and BRD4 normalized intensities for transcript amounts (**A**) and protein levels (**B**) across CCLE cancer lineages. For breast cancer, different colors indicate different breast cancer subtypes as indicated in the panel legend. (**C**) LOXL2-BRD4 protein correlation in TCGA-BRCA mass spectrometry study. Different colors indicate different breast cancer subtypes as indicated in the panel legend. (**D**) Cell viability of high and low LOXL2-expressing CCLE cell lines (protein levels) treated with different BETi small molecules at the indicated concentrations. The asterisks indicate the significance at the highest concentration using an unpaired Student's t-test. (*$P < 0.05$; ** $P < 0.005$; *** $P < 0.001$; **** $P < 0.0001$).

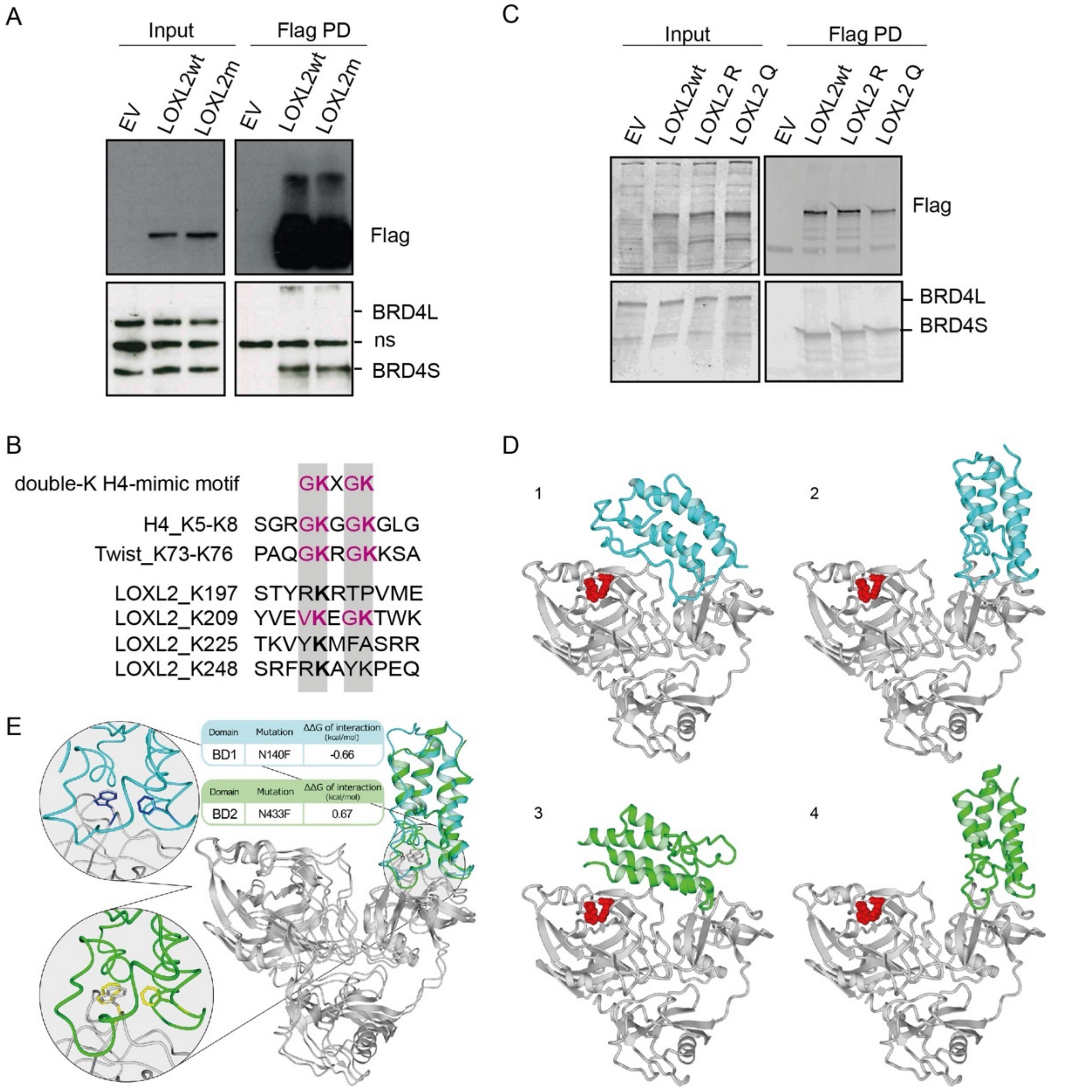
A
Input
Flag PD
EV
LOXL2wt
LOXL2m
EV
LOXL2wt
LOXL2m
Flag
BRD4L
ns
BRD4S
C
Input
Flag PD
EV
LOXL2wt
LOXL2 R
LOXL2 Q
EV
LOXL2wt
LOXL2 R
LOXL2 Q
Flag
BRD4L
BRD4S
B
double-K H4-mimic motif GKXGK
H4_K5-K8 SGRGKGGKGLG
Twist_K73-K76 PAQGKRGKKSA
LOXL2_K197 STYRKRTPVME
LOXL2_K209 YVEVKEGKTWK
LOXL2_K225 TKVYKMFASRR
LOXL2_K248 SRFRKAYKPEQ
D
1
2
3
4
E
Domain
Mutation
ΔΔG of interaction (kcal/mol)
BD1
N140F
-0.66
BD2
N433F
0.67

**Fig. S4. LOXL2-BRD4S interaction does not involve the activity of the bromodomains (A)** Flag PD in HEK293T cells overexpressing the empty vector (EV), LOXL2-Flag wild type (LOXL2wt), or the catalytically dead form of LOXL2-Flag (LOXL2m). Precipitates were analyzed by Western blot with the indicated antibodies. Three biological replicates were performed (ns: non-specific). (**B)** Schematic representation of the double-K H4-mimic motif partially shared between H4, Twist, and LOXL2. (**C)** Flag PD of MDA-MB-231 cells overexpressing EV, LOXL2wt, LOXL2 K209 residue mutated to R or LOXL2 K209-K212 residue mutated to Q. Precipitates were analyzed by western blot with the indicated antibodies. Three biological replicates were performed. (**D**) View of the 4 selected models highlighting LOXL2 histidines 626 and 628 (in red) whose mutations to glutamine showed not to affect the binding with BRD4. LOXL2 is represented in gray while BD1 and BD2 docking models in cyan and green, respectively. Panels 1, 2, 3, 4 correspond to docking models 4uyd_complex_10, 4uyd_complex_3, 2ouo_complex_4 and 2ouo_complex_5, respectively (**table S3**). (**E)** Details of docking models 4uyd_complex_3 (BD1), 2ouo_complex_5 (BD2), and their Asp→Phe mutant versions. The panel shows the superposition of LOXL2 (gray) docked to BD1 (cyan) and BD2 (green) with asparagines 140 and 433 mutated to phenylalanine (blue and yellow, respectively), both facing the buried triptofans 493 from LOXL2. On the left, zoomed visions of the mutants superposed over their wildtype structures.

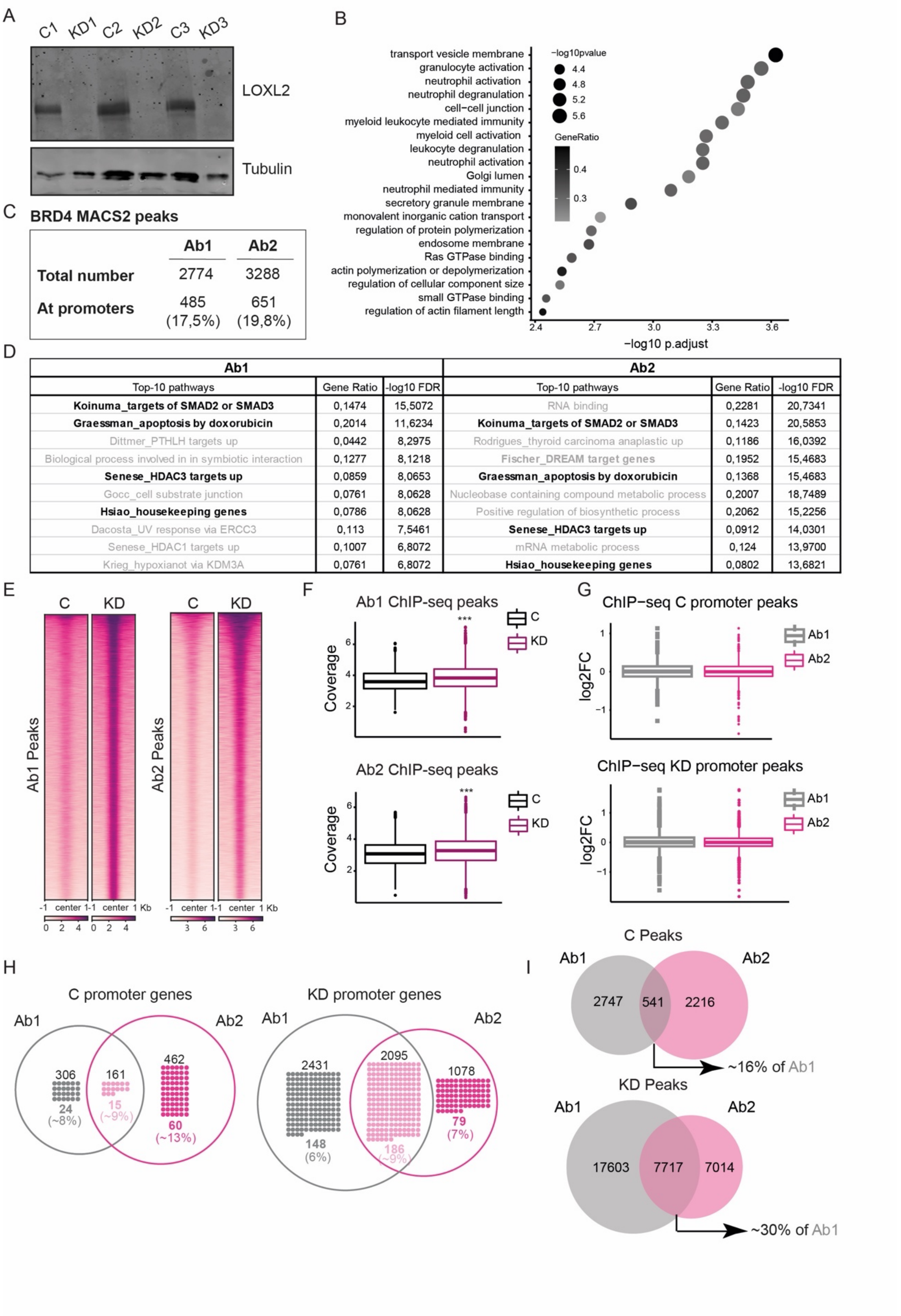

| Ab1 | | | Ab2 | | |
|---|---|---|---|---|---|
| Top-10 pathways | Gene Ratio | -log10 FDR | Top-10 pathways | Gene Ratio | -log10 FDR |
| **Koinuma_targets of SMAD2 or SMAD3** | 0,1474 | 15,5072 | RNA binding | 0,2281 | 20,7341 |
| **Graessman_apoptosis by doxorubicin** | 0,2014 | 11,6234 | **Koinuma_targets of SMAD2 or SMAD3** | 0,1423 | 20,5853 |
| Dittmer_PTHLH targets up | 0,0442 | 8,2975 | Rodrigues_thyroid carcinoma anaplastic up | 0,1186 | 16,0392 |
| Biological process involved in in symbiotic interaction | 0,1277 | 8,1218 | **Fischer_DREAM target genes** | 0,1952 | 15,4683 |
| **Senese_HDAC3 targets up** | 0,0859 | 8,0653 | **Graessman_apoptosis by doxorubicin** | 0,1368 | 15,4683 |
| Gocc_cell substrate junction | 0,0761 | 8,0628 | Nucleobase containing compound metabolic process | 0,2007 | 18,7489 |
| **Hsiao_housekeeping genes** | 0,0786 | 8,0628 | Positive regulation of biosynthetic process | 0,2062 | 15,2256 |
| Dacosta_UV response via ERCC3 | 0,113 | 7,5461 | **Senese_HDAC3 targets up** | 0,0912 | 14,0301 |
| Senese_HDAC1 targets up | 0,1007 | 6,8072 | mRNA metabolic process | 0,124 | 13,9700 |
| Krieg_hypoxianot via KDM3A | 0,0761 | 6,8072 | **Hsiao_housekeeping genes** | 0,0802 | 13,6821 |

**Fig. S5. ChIP-seq, RNA-seq, and ATAC-seq analysis of LOXL2 downregulation in MDA-MB-231 TNBC cells (A)** Western blot analysis of MDA-MB-231 cells infected with shControl (C) or shLOXL2 (KD) showing LOXL2 levels; Tubulin was used as a loading control. (**B)** Gene Set Enrichment Analysis (GSEA) of the genes up-regulated upon KD in the RNAseq dataset. (**C)** Number of BRD4 ChIP-seq total or promoter peaks using Ab1 or Ab2 antibodies identified with MACS2. (**D)** Top-10 GO-terms identified either with Ab1 or Ab2 when analyzing promoter peaks with the MSigDB. Gene Ratios and adjusted p-values are reported on the left side of each GS. In bold are GSs shared among the top-10 of Ab1 and Ab2. (**E)** Heatmap of ChIP-seq normalized signal (Reads per Genomic Content) in all peaks in C and KD for the antibodies Ab1 and Ab2. The normalized signal is calculated for a region of -1kb to 1kb from the center of the peaks. **(F)** Normalized ATAC-seq signal (Reads per Genomic Content) in the ChIP-seq peaks for Ab1 and Ab2 in shControl and shLOXL2 conditions. Significance was calculated using a two-sample Kolmogorov-Smirnov test. **(G)** RNA-seq logFC for genes associated with the peaks of Ab1 and Ab2 which fall in promoter regions in shControl and shLOXL2 respectively**.** Significance was calculated using a two-sample Kolmogorov-Smirnov test. (**H)** Venn diagram showing the number of promoter genes identified with the ChIP-seq with either Ab1 or Ab2 antibodies in C (left) or KD (right) conditions. DREAM target genes identified in each condition are depicted in colored dots. The numbers on top of the dots represent the total number of promoters retrieved for each condition. The numbers below the dots are respectively the (upper) total number of DREAM target gene promoters retrieved in each condition and (lower) the condition-relative percentage of DREAM target gene promoters identified (DREAM target gene promoters relative to all promoters). (**I)** Venn diagram showing the overlap between the total number of peaks detected with the ChIP-seq with either Ab1 or Ab2 in C (top) or KD (bottom) conditions. The overlap of Ab1 and Ab2 peaks is shown as a percentage (intersection relative to Ab1 peaks).

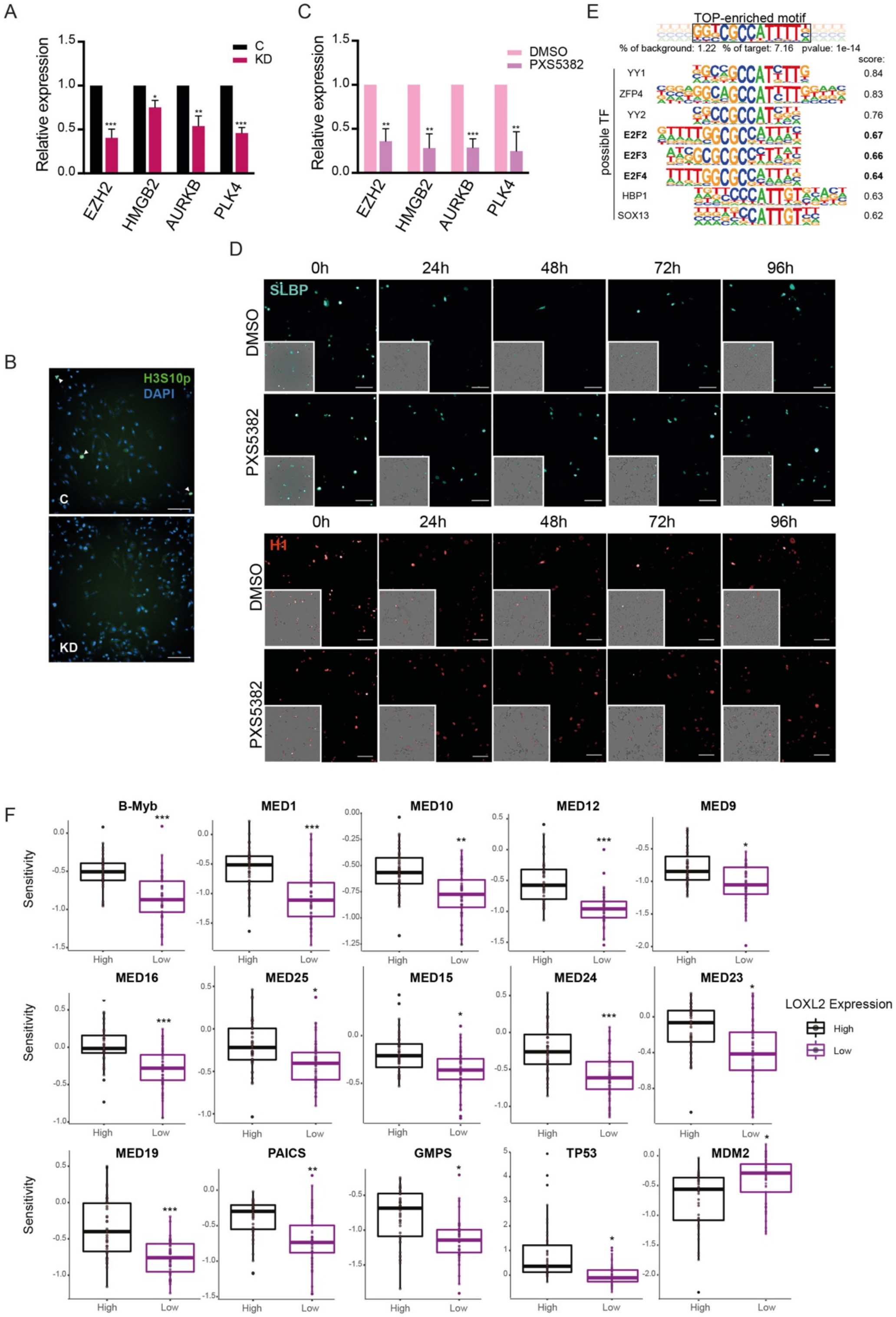
A
Relative expression
C
KD
EZH2
HMGB2
AURKB
PLK4
C
Relative expression
DMSO
PXS5382
EZH2
HMGB2
AURKB
PLK4
E
TOP-enriched motif
% of background: 1.22 % of target: 7.16 pvalue: 1e-14
score:
possible TF
YY1 0.84
ZFP4 0.83
YY2 0.76
E2F2 0.67
E2F3 0.66
E2F4 0.64
HBP1 0.63
SOX13 0.62
B
H3S10p
DAPI
C
KD
D
0h
24h
48h
72h
96h
SLBP
DMSO
PXS5382
H1
F
B-Myb
MED1
MED10
MED12
MED9
MED16
MED25
MED15
MED24
MED23
MED19
PAICS
GMPS
TP53
MDM2
Sensitivity
High
Low
LOXL2 Expression

**Fig. S6. LOXL2 repression affects the transcriptional regulation of DREAM target genes (A)** Real-time quantitative PCR (qPCR) showing the changes in mRNA expression of four selected DREAM target genes (EZH2, HMGB2, AURKB, and PLK4) in MDA-MB-231 cells treated with shControl (C) or shLOXL2 (KD). Gene expression was normalized against an endogenous control (Pumilio homolog 1) and represented as the expression relative to the C condition, which was set as 1. The asterisks indicate the significance using a paired Student's t-test (*$P < 0.05$; ** $P < 0.005$; *** $P < 0.001$). (**B)** Representative images of H3S10P high throughput immunofluorescence in MDA-MB-231 treated either with C or KD. Scale bar; 100µm. (**C)** Real-time quantitative PCR (qPCR) showing the changes in mRNA expression of four selected DREAM target genes (EZH2, HMGB2, AURKB, and PLK4) in MDA-MB-231 cells treated with DMSO or PXS5382. Gene expression was normalized against an endogenous control (Pumilio homolog 1) and represented as the expression relative to the DMSO condition, which was set as 1. The asterisks indicate the significance using a paired Student's t-test (*$P < 0.05$; ** $P < 0.005$; *** $P < 0.001$). (**D)** MDA-MB-231 cells expressing SLBP-mTurquoise2 and H1-Maroon1 treated with DMSO or PXS5382 for 96h. Representative images of the quantification in **Fig. 4D** are shown. Images of big panels show SLBP-mTurquoise2 (top) and H1-Maroon1 (bottom) while small panels show their overlap with brightfield. Scale bar; 100µm. **(E)** Scores of possible transcription factors binding to the top-enriched transcription factor binding motif retrieved with the Homer-*de novo* analysis. **(F)** Gene-specific differential gene essentiality between high and low LOXL2-expressing cell lines (CCLE) as calculated analyzing the Achilles' dataset. Significance was determined using the Student's t-test with BH multiple hypothesis correction (*$P < 0.05$; ** $P < 0.005$; *** $P < 0.001$).

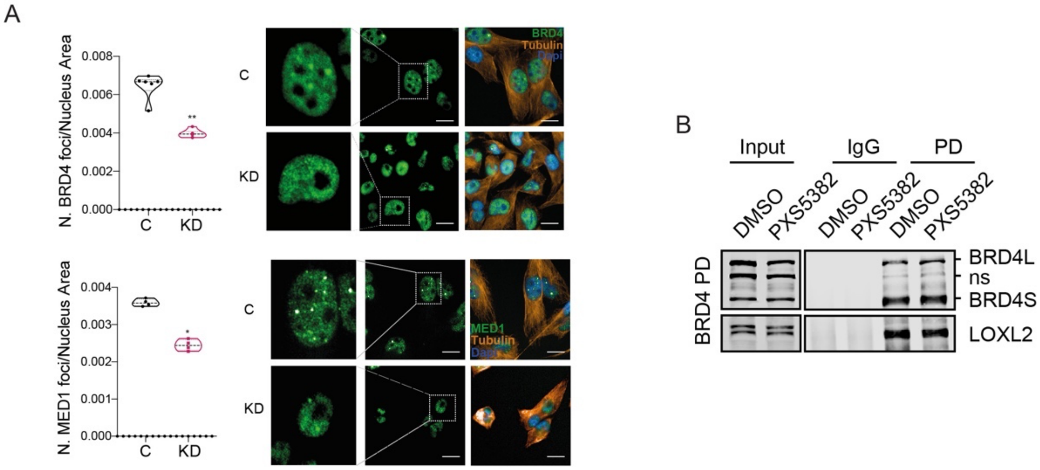
A
N. BRD4 foci/Nucleus Area
0.008
0.006
0.004
0.002
0.000
C
KD
**
BRD4
Tubulin
Dapi
N. MED1 foci/Nucleus Area
0.004
0.003
0.002
0.001
0.000
*
MED1
B
Input
IgG
PD
DMSO
PXS5382
BRD4 PD
BRD4L
ns
BRD4S
LOXL2

**Fig. S7**. **LOXL2 inhibition impairs the formation of BRD4 and MED1 nuclear foci (A)** High-throughput immunofluorescence of BRD4 (top) and MED1 (bottom) in shControl (C) or shLOXL2 (KD) infected MDA-MB-231 cells. Quantification of foci number corrected by nucleus area (left). Representative images showing BRD4 or MED1 nuclear foci (green); Tubulin (orange) and DAPI (blue) were used as cytoplasm and nuclear markers, respectively (right). Scale bar; 10µm. (**B)** BRD4 PD in MDA-MB-231 cells treated with DMSO or PXS5382 for 96h. Precipitates were analyzed by Western blot with the indicated antibodies. Irrelevant IgGs were used as a negative control (ns: non-specific). Three biological replicates were performed.

| model ID | residues | source | domain |
|---|---|---|---|
| 4uyd | 44-183 | PDB | BR1 |
| 2ouo | 333-460 | PDB | BR2 |
| AF-O60885-F1 | 1-1362 | Uniprot | FULL |
| O60885-O60885-EXP-5khm | 44-168 | Interactome3d | BR1 |
| O60885-O60885-EXP-6djc | 44-173 | Interactome3d | BR1 |
| O60885-O60885-EXP-6g0s | 42-168 | Interactome3d | BR1 |
| O60885-P40337-EXP-5t35 | 333-460 | Interactome3d | BR2 |
| O60885-P40337-EXP-6sis | 333-460 | Interactome3d | BR2 |
| O60885-Q04206-EXP-4kv1 | 41-168 | Interactome3d | BR1 |

**Table S1. BRD4 structures** List of BRD4 structures used in the analysis, showing covered sequence, data source and covered domain.

| BD1 | | | | | |
|---|---|---|---|---|---|
| model ID | residues | Interaction (kcal/mol) | Stability (kcal/mol) | surface (A^2) | software |
| docking results | | | | | |
| 4uyd_zdock_10 | 44-183 | -15,86 | 31,00 | 853,04 | zdock |
| 4uyd_zdock_2 | 44-183 | -14,55 | 24,45 | 686,69 | zdock |
| 4uyd_zdock_5 | 44-183 | -13,71 | 25,83 | 681,08 | zdock |
| 4uyd_zdock_3 | 44-183 | -13,14 | 28,77 | 835,23 | zdock |
| 4uyd_zdock_1 | 44-183 | -11,07 | 25,80 | 634,88 | zdock |
| 4uyd_zdock_4 | 44-183 | -9,86 | 23,64 | 648,02 | zdock |
| O60885-O60885-EXP-5khm_5ze3_0 | 44-168 | -9,76 | -31,14 | 849,73 | fishing |
| 4uyd_zdock_8 | 44-183 | -9,38 | 28,24 | 555,69 | zdock |
| 4uyd_zdock_7 | 44-183 | -9,18 | 20,34 | 639,86 | zdock |
| O60885-O60885-EXP-5khm_5ze3_1 | 44-169 | -9,15 | -23,47 | 629,34 | fishing |
| 4uyd_zdock_6 | 44-183 | -8,08 | 29,10 | 761,08 | zdock |
| O60885-O60885-EXP-6g0s_5ze3_0 | 44-167 | -7,20 | -26,29 | 443,91 | fishing |
| O60885-Q04206-EXP-4kv1_5ze3_0 | 41-168 | -6,94 | -28,39 | 890,63 | fishing |
| O60885-O60885-EXP-6djc_5ze3_3 | 42-172 | -6,79 | -29,84 | 398,57 | fishing |
| 4uyd_zdock_9 | 44-183 | -5,18 | 38,56 | 450,44 | zdock |
| O60885-O60885-EXP-5khm_5ze3_2 | 44-170 | -4,38 | -26,07 | 271,28 | fishing |
| 4uyd_vina | 44-183 | -3,88 | 33,10 | 405,63 | vina |
| O60885-O60885-EXP-6djc_5ze3_1 | 42-172 | -3,29 | -26,05 | 234,31 | fishing |
| O60885-O60885-EXP-6g0s_5ze3_5 | 44-167 | -2,47 | -25,28 | 141,86 | fishing |
| O60885-O60885-EXP-6djc_5ze3_2 | 42-172 | -1,91 | -31,04 | 231,78 | fishing |
| O60885-O60885-EXP-6djc_5ze3_0 | 42-172 | -0,96 | -27,86 | 116,97 | fishing |
| O60885-O60885-EXP-5khm_5ze3_3 | 44-171 | -0,86 | -24,77 | 213,54 | fishing |
| O60885-O60885-EXP-6g0s_5ze3_2 | 44-167 | -0,45 | -25,29 | 113,96 | fishing |
| O60885-O60885-EXP-6g0s_5ze3_1 | 44-167 | 0,27 | -26,18 | 69,58 | fishing |
| top 10 results | | | | | |
| 4uyd_zdock_10 | 44-183 | -15,86 | 31,00 | 853,04 | zdock |
| 4uyd_zdock_2 | 44-183 | -14,55 | 24,45 | 686,69 | zdock |
| 4uyd_zdock_5 | 44-183 | -13,71 | 25,83 | 681,08 | zdock |
| 4uyd_zdock_3 | 44-183 | -13,14 | 28,77 | 835,23 | zdock |
| 4uyd_zdock_1 | 44-183 | -11,07 | 25,80 | 634,88 | zdock |
| 4uyd_zdock_4 | 44-183 | -9,86 | 23,64 | 648,02 | zdock |
| O60885-O60885-EXP-5khm_5ze3_0 | 44-168 | -9,76 | -31,14 | 849,73 | fishing |
| 4uyd_zdock_8 | 44-183 | -9,38 | 28,24 | 555,69 | zdock |
| 4uyd_zdock_7 | 44-183 | -9,18 | 20,34 | 639,86 | zdock |
| O60885-O60885-EXP-5khm_5ze3_1 | 44-169 | -9,15 | -23,47 | 629,34 | fishing |
| filtered by clustering and agreement with alphafold model | | | | | |
| 4uyd_zdock_10 | 44-183 | -15,86 | 31,00 | 853,04 | zdock |
| 4uyd_zdock_4 | 44-183 | -9,86 | 23,64 | 648,02 | zdock |
| 4uyd_zdock_3 | 44-183 | -13,14 | 28,77 | 835,23 | zdock |
| O60885-O60885-EXP-5khm_5ze3_1 | 44-169 | -9,15 | -23,47 | 629,34 | fishing |
| filtered by residue incompatibility and manual curation | | | | | |
| 4uyd_zdock_10 | 44-183 | -15,86 | 31,00 | 853,04 | zdock |
| 4uyd_zdock_3 | 44-183 | -13,14 | 28,77 | 835,23 | zdock |

| BD2 | | | | | |
|---|---|---|---|---|---|
| model ID | residues | Interaction (kcal/mol) | Stability (kcal/mol) | surface (A^2) | software |
| docking results | | | | | |
| 2ouo_zdock_8 | 333-460 | -24,54 | 19,87 | 1322,34 | zdock |
| 2ouo_zdock_4 | 333-460 | -15,12 | 28,49 | 733,17 | zdock |
| 2ouo_zdock_3 | 333-460 | -14,31 | 36,58 | 1082,26 | zdock |
| 2ouo_zdock_9 | 333-460 | -14,29 | 33,01 | 927,98 | zdock |
| 2ouo_zdock_5 | 333-460 | -12,32 | 19,27 | 639,85 | zdock |
| 2ouo_zdock_7 | 333-460 | -11,91 | 39,07 | 832,66 | zdock |
| 2ouo_zdock_1 | 333-460 | -11,42 | 37,57 | 844,41 | zdock |
| 2ouo_zdock_2 | 333-460 | -11,00 | 20,45 | 611,46 | zdock |
| 2ouo_zdock_6 | 333-460 | -7,80 | 30,73 | 829,39 | zdock |
| 2ouo_zdock_10 | 333-460 | -7,25 | 29,40 | 739,58 | zdock |
| 2ouo_vina | 333-460 | -3,40 | 45,80 | 585,29 | vina |
| O60885-P40337-EXP-5t35_5ze3_1 | 349-459 | -2,45 | -27,45 | 203,52 | fishing |
| O60885-P40337-EXP-6sis_5ze3_0 | 349-459 | -2,26 | -23,80 | 139,74 | fishing |
| O60885-P40337-EXP-6sis_5ze3_1 | 349-459 | -1,65 | -26,44 | 194,26 | fishing |
| O60885-P40337-EXP-5t35_5ze3_4 | 349-459 | -1,39 | -28,46 | 146,32 | fishing |
| top 10 results | | | | | |
| 2ouo_zdock_8 | 333-460 | -24,54 | 19,87 | 1322,34 | zdock |
| 2ouo_zdock_4 | 333-460 | -15,12 | 28,49 | 733,17 | zdock |
| 2ouo_zdock_3 | 333-460 | -14,31 | 36,58 | 1082,26 | zdock |
| 2ouo_zdock_9 | 333-460 | -14,29 | 33,01 | 927,98 | zdock |
| 2ouo_zdock_5 | 333-460 | -12,32 | 19,27 | 639,85 | zdock |
| 2ouo_zdock_7 | 333-460 | -11,91 | 39,07 | 832,66 | zdock |
| 2ouo_zdock_1 | 333-460 | -11,42 | 37,57 | 844,41 | zdock |
| 2ouo_zdock_2 | 333-460 | -11,00 | 20,45 | 611,46 | zdock |
| 2ouo_zdock_6 | 333-460 | -7,80 | 30,73 | 829,39 | zdock |
| 2ouo_zdock_10 | 333-460 | -7,25 | 29,40 | 739,58 | zdock |
| filtered by clustering and agreement with alphafold model | | | | | |
| 2ouo_zdock_8 | 333-460 | -24,54 | 19,87 | 1322,34 | zdock |
| 2ouo_zdock_4 | 333-460 | -15,12 | 28,49 | 733,17 | zdock |
| 2ouo_zdock_5 | 333-460 | -12,32 | 19,27 | 639,85 | zdock |
| filtered by residue incompatibility and manual curation | | | | | |
| 2ouo_zdock_4 | 333-460 | -15,12 | 28,49 | 733,17 | zdock |
| 2ouo_zdock_5 | 333-460 | -12,32 | 19,27 | 639,85 | zdock |

**Table S2. Possible BRD4-LOXL2 interaction models** List and statistics of generated docking models. Results are divided by BD1 and BD2 and for each model are reported: BRD4 covered sequence, FoldX interaction energy, and stability in kcal/mol, buried surface in Å2. The table progresses downwards showing how candidate complexes have been filtered to finish with the four final models.

| BD1 | | | |
|---|---|---|---|
| mutationin model | mutation's molecule | DDG of interaction | invalidating interaction |
| O60885-O60885-EXP-5khm_5ze3_1 | | | |
| W81A | BRD4 | 1,91 | F |
| 4uyd_zdock_3 | | | |
| R447A | LOXL2 | 3,69 | F |
| W493A | LOXL2 | 1,39 | F |
| Y494A | LOXL2 | 1,71 | F |
| Y537A | LOXL2 | 3,44 | F |
| L92A | BRD4 | 1,60 | F |
| D144A | BRD4 | -2,17 | T |
| D145A | BRD4 | -1,61 | F |
| N140F | BRD4 | -0,66 | F |
| 4uyd_zdock_4 | | | |
| W493A | LOXL2 | 3,88 | F |
| Y537A | LOXL2 | 1,75 | F |
| L94A | BRD4 | 1,77 | F |
| N140F | BRD4 | 3,24 | T |
| 4uyd_zdock_10 | | | |
| E445A | LOXL2 | 1,45 | F |
| W493A | LOXL2 | 2,67 | F |
| W622A | LOXL2 | 1,50 | F |
| Y629A | LOXL2 | 2,05 | F |
| P46A | BRD4 | 1,58 | F |
| R58A | BRD4 | 1,93 | F |
| K102A | BRD4 | 2,43 | F |
| W120A | BRD4 | 2,67 | F |

| BD2 | | | |
|---|---|---|---|
| mutationin model | mutation's molecule | DDG of interaction | invalidating interaction |
| 2ouo_zdock_4 | | | |
| R423A | BRD4 | 1,51 | F |
| Y432A | BRD4 | 1,79 | F |
| P435A | BRD4 | 1,74 | F |
| W493A | LOXL2 | 1,73 | F |
| Y494A | LOXL2 | 1,98 | F |
| Y537A | LOXL2 | 1,60 | F |
| H618A | LOXL2 | 1,77 | F |
| Y629A | LOXL2 | 1,80 | F |
| F676A | LOXL2 | 1,68 | F |
| N433F | BRD4 | -0,07 | F |
| 2ouo_zdock_5 | | | |
| P375A | BRD4 | 1,80 | F |
| L387A | BRD4 | 1,41 | F |
| E438A | BRD4 | 1,35 | F |
| R447A | LOXL2 | 3,14 | F |
| W493A | LOXL2 | 2,77 | F |
| Y537A | LOXL2 | 2,84 | F |
| N433F | BRD4 | 0,67 | F |
| 2ouo_zdock_8 | | | |
| W374A | BRD4 | 2,84 | F |
| L387A | BRD4 | 1,49 | F |
| Y390A | BRD4 | 1,34 | F |
| D399A | BRD4 | 1,69 | F |
| R441A | LOXL2 | 1,42 | F |
| R447A | LOXL2 | 2,75 | F |
| F489A | LOXL2 | 1,67 | F |
| W493A | LOXL2 | 3,35 | F |
| Y494A | LOXL2 | 2,62 | F |
| W495A | LOXL2 | 1,63 | F |
| Y537A | LOXL2 | 1,53 | F |
| N433F | BRD4 | 4,20 | T |

**Table S3**. **In silico mutagenesis residues** In silico mutation analysis for the top 7 docking models. Results are divided by BD1 and BD2 and for each model are reported: interface residues which mutated to alanine shows a relevant change in interaction energy, molecule to which the interface residue belongs, predicted ΔΔG of interaction for the mutation, whether the predicted effect can invalidate the model. For each model implying asparagines 140 (BD1) or 433 (BD2) in the interaction, we show (bold box) the effect of mutating to phenylalanine.

Homer *de novo* Motif Results (excluding possible false positive)

| Enriched Motifs | P-value | log P-pvalue | % of Targets | % of Background |
|---|---|---|---|---|
| 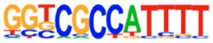 | 1E-14 | -3,34E+04 | 7.16% | 1.22% |
| 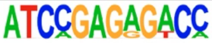 | 1E-12 | -2,818E+04 | 1.12% | 0.00% |
| 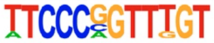 | 1E-12 | -2,808E+04 | 1.34% | 0.01% |
| 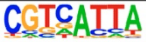 | 1E-12 | -2,765E+04 | 24.16% | 12.02% |

**Table S4. Transcription factor motif analysis** Statistically significant transcription factor motif analysis of Ab1 promoter peaks retrieved with HOMER motif analysis software.

**Movie S1**. (Separate file)

**Time-lapse of mTurquoise-SLBP expressing cells** Representative time-lapse movie of MDA-MB-231 cells expressing mTurquoise2-SLBP and H1-Maroon1, treated with DMSO or PXS5382 for 96h. Images were acquired every 15 minutes.
